# Injury distance limits the transcriptional response to spinal injury

**DOI:** 10.1101/2024.05.27.596075

**Authors:** Zimei Wang, Manojkumar Kumaran, Elizabeth Batsel, Sofia Testor-Cabrera, Zac Beine, Alicia Alvarez Ribelles, Pantelis Tsoulfas, Ishwariya Venkatesh, Murray G. Blackmore

## Abstract

The ability of neurons to sense and respond to damage is fundamental to homeostasis and nervous system repair. For some cell types, notably dorsal root ganglia (DRG) and retinal ganglion cells (RGCs), extensive profiling has revealed a large transcriptional response to axon injury that determines survival and regenerative outcomes. In contrast, the injury response of most supraspinal cell types, whose limited regeneration constrains recovery from spinal injury, is mostly unknown. Here we employed single-nuclei sequencing in mice to profile the transcriptional responses of diverse supraspinal cell types to spinal injury. Surprisingly, thoracic spinal injury triggered only modest changes in gene expression across all populations, including corticospinal tract (CST) neurons. Moreover, CST neurons also responded minimally to cervical injury but much more strongly to intracortical axotomy, including upregulation of numerous regeneration and apoptosis-related transcripts shared with injured DRG and RGC neurons. Thus, the muted response of CST neuron to spinal injury is linked to the injury’s distal location, rather than intrinsic cellular characteristics. More broadly, these findings indicate that a central challenge for enhancing regeneration after a spinal injury is the limited sensing of distant injuries and the subsequent modest baseline neuronal response.

## Introduction

Axon injury can trigger diverse responses in neurons, ranging from rapid cell death, cell survival with abortive axonal sprouting, or robust axonal regeneration^1–5^. Underlying these differences are distinct patterns of gene transcription that are triggered as neurons sense and respond to axon damage^6–8^. Clarifying the various transcriptional responses to injury is a powerful means to develop strategies to protect vulnerable cell types and to identify both pro-and anti-growth pathways that can be targeted to spur regeneration. These efforts rely on a detailed and accurate description of the baseline injury response and thus a major effort in regenerative neuroscience has been to profile the transcriptional reaction of neurons to axon damage^9–15^

For some cell types, notably sensory neurons of the dorsal root ganglia (DRG)^10,15–17^ and retinal ganglion cells (RGCs)^11,12,18,19^, numerous profiling efforts have provided deep insight. In both cell types, work using bulk and single-cell or single-nuclei sequencing has detected many hundreds to thousands of strongly differentially expressed genes (DEGs) after injury ^10–12,16,18^. DRG neurons, which survive axotomy and mount an effective regenerative response, respond to injury with a partial loss of transcripts that distinguish subtypes and the upregulation of numerous transcripts that support axon growth. RGCs are highly vulnerable to cell death and display limited spontaneous axon growth after axotomy but can be coaxed to regenerate long axons using a variety of interventions ^20–25^. Accordingly, the transcriptional response of RGCs shows strong upregulation of transcripts associated with both cell death and axon regeneration. Consequently, experimental interventions that dampen the damage detection pathways confer neuroprotection but interfere with spontaneous and treatment-induced axon regeneration^2,19,26^.

In contrast to DRG and RGC neurons, very little is known about the genome-wide responses of supraspinal neurons after spinal injury. Phenotypically, most supraspinal populations survive spinal axotomy but fail to initiate long-distance axon regeneration, although some spontaneously grow collateral branches above the injury^1,27^. For most populations the transcriptional basis for the low-growth response remains largely unknown. Conceptually, it is unclear whether axon extension is constrained by an inability to activate necessary pro-growth genes or if these genes are expressed but countered by emerging growth-suppressive networks. The corticospinal tract (CST), an important regulator of sensation and fine motor control ^28–30^, is among the most studied descending tracts and follows the pattern of spontaneous collateralization but not extension of the main axon^1,31^. Earlier studies that examined specific transcripts in supraspinal populations, including the CST, showed minimal transcriptional responses to injury, particularly when axotomy occurred far from the cell body^32,33^. In contrast, recent profiling of the CST after spinal injury reported a large-scale transcriptional response and a transient reversion to an embryonic state^9^. Given the importance of supraspinal populations to restoring function after spinal injuries it is critical to expand understanding of the response from less studied supraspinal cell types and to clarify the response of CST.

In this study we utilized single-nuclei sequencing techniques in a mouse model to reveal how adult supraspinal projection neurons respond transcriptionally to spinal injury. After thoracic injury we detected only modest transcriptional changes in any supraspinal population, including the CST, with DEGs numbering fewer than 10% of the totals typically reported for DRG and RGC neurons. To explore the effect of cell-body-to-axotomy distance we focused on the CST and performed additional profiling experiments after injury in the cervical spinal cord or within the cortex, very near to CST cell bodies. Intracortical injury, unlike cervical injury, triggered transcriptional changes on a scale similar to those previously observed in DRG and RGC responses. Moreover, intersection of the CST intracortical response with those of DRG and RGC identified a core network of injury-responsive genes, many of which were previously linked to apoptosis and/or axon regeneration. Remarkably, CSTs that are injured in the spinal region showed minimal activation of this network. These data provide comprehensive insight into the response of supraspinal neurons to spinal injury and favor a model in which limited damage response, and not counteracting transcription or a failure to maintain an initial reversion to a growth state, likely explains the limited axon growth after spinal injury.

## Results

### Profiling supraspinal transcriptional responses to thoracic injury

We used retrograde labeling, cell nuclei isolation, and single-nuclei analyses to examine gene expression in supraspinal populations after spinal injury. Cell nuclei of supraspinal neurons were fluorescently labeled by injection of AAV2-retro-H2B-mGl to the lumbar spinal cord of adult mice, which we previously found to produce bright, nuclear-localized signal in tens of thousands of supraspinal neurons throughout the brain^34,35^ (**Fig. 1a**). One week later, following successful viral transduction of supraspinal neurons, we performed a complete crush of the spinal cord at thoracic level 10 (T10)). After another week, fluorescently labeled brain regions were micro-dissected and flash frozen. Supraspinal nuclei were purified by fluorescence-activated nuclei sorting (FANS) (**Sup. Fig. 1**), and single-nuclei libraries prepared on a 10X Genomics platform with three animals pooled in each of three replicate samples. Spinal tissue was collected at the time of dissection and injury completeness was confirmed in spinal sections subjected to GFAP immunohistochemistry (**Fig. 1a, inset**).

**Figure 1:**
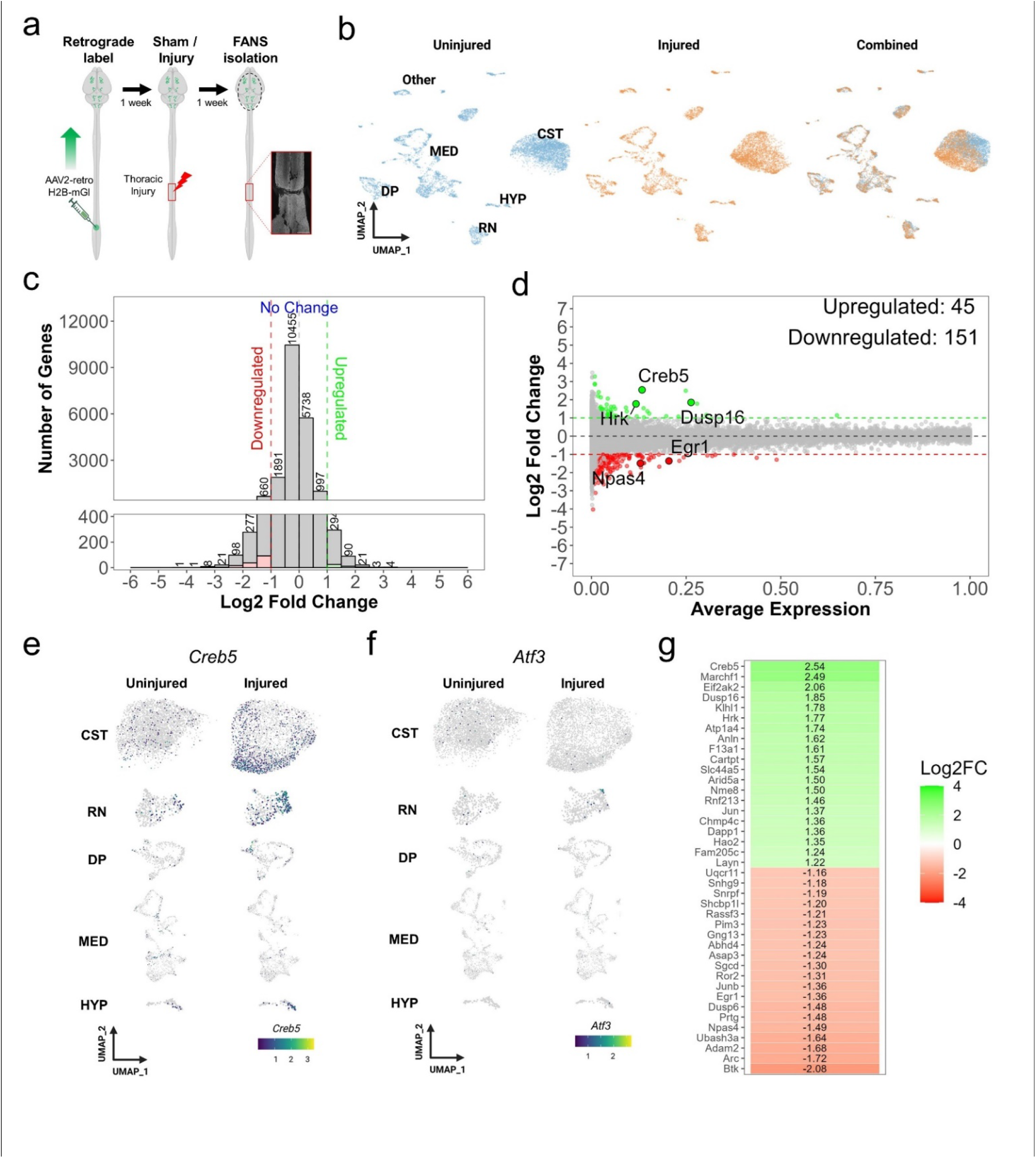
Single-nuclei analyses detect minimal changes in gene expression in supraspinal neurons following thoracic spinal injury. (a) Experimental design including retrograde labeling of supraspinal neurons, thoracic spinal injury, and fluorescent activated nuclei sorting one week post-injury. Inset shows GFAP verification of spinal injury. (b) UMAP visualization of uninjured (blue) and injured (orange) nuclei, both of which cluster into six main cell types: CST (Corticospinal Tract), DP (Dorsal Pons), RN (Red Nucleus neurons), MED (Medulla neurons), HYP (Hypothalamus neurons), and ‘Other’ cell types (c) A histogram indicating the number of Differentially Expressed Genes in injured versus uninjured CST neurons. (d) MA plot showing the relationship between transcript abundance and differential gene expression in injured versus uninjured CST neuros, with significantly up- and down-regulation indicated by green and red, respectively (non-parametric Wilcoxon rank sum test, p-value<0.05). (f) *Crebs* and (g) *Atf3* gene expression across five cell types under uninjured and injured states, indicating varying responses to injury (h) A heatmap indicating the identity of the top twenty up- and down regulated transcripts in injured CST neurons.

To identify injury-sensitive transcripts we used a Seurat-based pipeline to merge thoracic-injured samples with uninjured control samples from our prior work^34^. The timing of viral delivery and methods of dissection, FANS purification, and library preparation were identical between uninjured and injured samples. UMAP clustering of the merged datasets revealed sixteen distinct clusters, which were assigned supraspinal cell identity based on our previously established markers ^34^(**Sup. Fig. 2a-c**). These clusters were then grouped into five main brain regions: CST (corticospinal), HYP (hypothalamus), RN (red nucleus), DP (dorsal pons) and MED (medullary neurons, predominantly reticular) (**Sup. Fig. 2c)**. Notably, injured cells were found in the same UMAP clusters as uninjured cells (**Fig. 1b**) and continued to express cell-specific transcripts (**Sup. Fig. 2d)**. For example, injured CST neurons clustered separately from all subcortical cell types, intermingled with uninjured CST neurons, and remained readily identifiable by markers such as Crym^36^ (**Sup. Fig.2c, d**). Thus, unlike in peripherally injured DRG neurons^10^, axotomy does not trigger widespread de-differentiation in supraspinal neurons.

Indeed, supraspinal neurons typically showed limited transcriptional responses to thoracic injury. When all nuclei were pooled for a global comparison between injured and uninjured neurons, only 147 transcripts changed significantly above a threshold of two-fold (log2 fold change ±1; 41 increasing, 106 decreasing) (**Sup. Fig. 3)**. Furthermore, nearly 90% of the 147 DEGs came from transcripts detected in fewer than 10% of all nuclei, thus minimizing the effect on the overall population. In contrast, many hundreds or thousands of transcripts typically surpass the same threshold following axotomy in other neuronal cell types (see **Fig. 6** below and ^10–12,16^). Comparisons performed population-by-population also revealed few injury-regulated transcripts (**Sup. Fig. 3**). Of all supraspinal types, CST neurons showed the largest number of transcripts significantly changed, with 45 and 151 up-and downregulated more than two-fold, respectively (**Fig. 1c,d**). As in the all-neuron analysis, nearly all the CST DEGs were low-abundance transcripts (**Fig. 1d**). Gene ontology (GO) enrichment analysis of up-and down-regulated gene sets revealed no over-represented terms, possibly due to the limited size of the input lists. Interestingly, *Creb5, Hrk*, and *Dusp16* emerged as some of the most upregulated transcripts, all of which have been previously associated with axon damage^26,37,38^ (**Fig. 1e, g**). On the other hand, other transcripts typical of axotomized or otherwise stressed cells, such as *Atf3, Ddit3, Ecel1*, and *Casp3* were not upregulated (**Fig, 1g**). Likewise, transcripts associated with neural differentiation, including *Sox2* and *Ascl1*^9^, remained unaffected by injury (all thoracic injury DEGs are available in **Supplemental Table. 1**). Examination of the list of downregulated transcripts showed several markers of neural activity (*Npas4, Arc*, and *Egr1*) but not transcripts linked to neural differentiation or to axon growth (**Fig. 1g, Supplemental Table. 1**). In summary, profiling following thoracic injury revealed a transcriptional response characterized by the activation of a limited set of damage-responsive transcripts and a decrease in several activity-dependent transcripts. However, there was not widespread activation of transcripts associated with cellular stress, embryonic development, or axon growth.

### The response of CST neurons to cervical axotomy

A potential explanation for the muted transcriptional response to thoracic injury could be the significant distance between the injury site and the originating cell bodies^32,33^. To test if a closer injury site elicits a more pronounced response, we moved the injury site to cervical spinal cord. This experiment focused exclusively on CST neurons, which were selected based on their abundance, functional importance, and the feasibility of transecting nearly all descending CST axons via dorsal hemisection while maintaining animal survival. Neurons were retrogradely labeled by C5 injection of AAV2-retro-H2B-mGl followed by unilateral dorsal hemisection at the C4/5 level. One week later layer V cortical tissue was microdissected and flash frozen, replicate libraries were prepared from FANS-purified CST nuclei from three pooled animals, and data from injured and uninjured libraries were merged and analyzed in Seurat. Spinal tissue from injured animals was imaged with GFAP immunohistochemistry to confirm complete transection of the right CST (**Fig. 2a, inset**). In an initial check of cell purity, small clusters were identified that expressed markers for superficial neurons (*Cux2*) or intracortically-projecting neurons (*Slc30a2*); these comprised 1.7% of total nuclei and were removed^34,39^. The remaining cluster contained a mixture of nuclei from injured and uninjured samples, both of which expressed similar levels of CST and layer V markers *Crym, Bcl11b*, and *Fezf2* (**Fig. 2b,c)**. These data indicate high purity of CST nuclei and, similarly to the thoracic injury, an absence of injury-triggered dedifferentiation that was reported in axotomized DRG neurons^10^. Consistent with this, only 32 DEGs exceeded a two-fold threshold (**Fig. 2d, e**). Although the list was small, it showed a highly significant overlap with the set of thoracic DEGs (46.8%, 32.2-fold enrichment, p < 7.409e-20, hypergeometric test) including shared upregulation of *Creb5, Hrk*, and *Dusp16* and shared downregulation of *Npas4, Arc, and Egr1* (**Fig. 2e,f**). This consistency across different injury locations enhances confidence in the snRNA-seq method’s ability to detect a shared biological signal, albeit small. However, even with axotomy moved to the cervical level, it still produced limited transcriptional changes in CST neurons.

**Figure 2:**
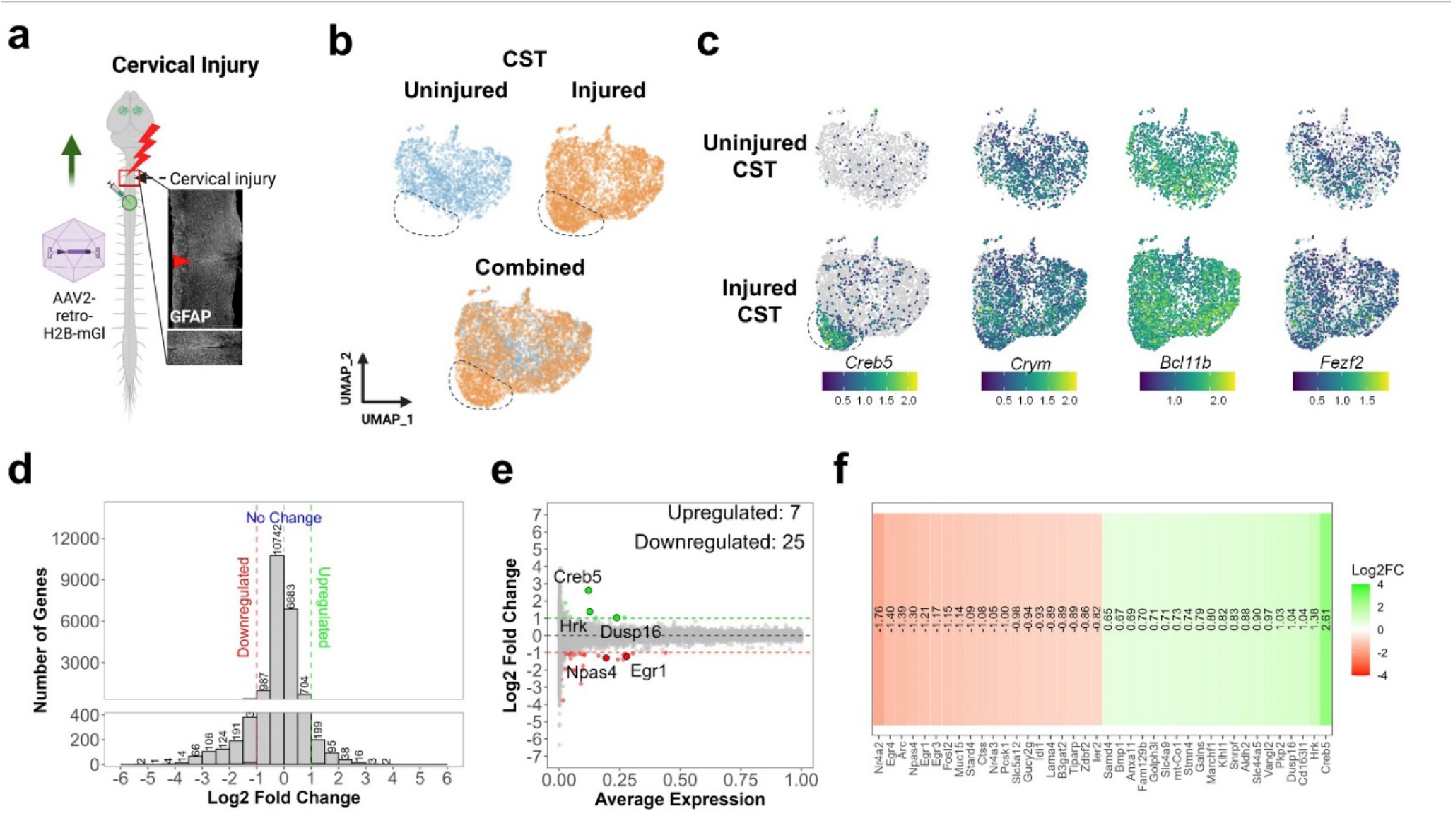
Single-nuclei profiling detects minimal responses in CST neurons to cervical spinal injury. (a) Experimental design: CST neurons were retrogradely labeled with nuclear-localized H2B-mGI, the right spinal cord was transected or left uninjured, and one week later nuclei were harvested for single-nuclei sequencing. Inset shows GFAP signal, verifying injury completeness, (b) UMAP visualization shows predominant overlap of injured and uninjured nuclei, with the exception of one injured-enriched group (dotted circle), (c) Feature plots show expression of *Crym, Bcl11b*, and *Fezf2*, verifying CST identity, and higher levels of *Creb*5 in the injured-enriched region (dotted line), (d) A histogram indicating the number of Differentially Expressed Genes in injured versus uninjured CST neurons, (e) MA plot showing the relationship between transcript abundance and differential gene expression in injured versus uninjured CST neuros. In both (d) and (e) transcripts that are significantly up- and down-regulated are colored green and red, respectively (non-parametric Wilcoxon rank sum test, p-value<0.05). (f) A heatmap indicating the identity of the top twenty up- and down-regulated transcripts in CST neurons one week after cervical spinal injury.

We next explored the idea that while CST neurons as a collective exhibit a modest response to injury, there might be subpopulations within them that show greater responsiveness. Indeed, although the injured and uninjured samples mostly intermingled in UMAP clustering, closer examination revealed a region of non-overlap that was comprised almost entirely of injured nuclei. Interestingly, expression of *Creb5*, previously associated with neuronal injury responses^37^, was highly concentrated in this region (**Fig. 2c, dotted circle**). When injured nuclei within this region were compared separately to uninjured neurons they displayed somewhat larger differences, with 210 transcripts exceeding a threshold of ±2-fold change and more than 551 at a relaxed threshold of ±50% (± 0.58 log2FC) (**Supplemental Table 2, Sup. Fig. 4**). We therefore designated injured nuclei within this region, which comprised 19.1% of all injured nuclei, as CST-high *Creb5* (CST-hCreb5). Among the genes significantly upregulated in this subgroup were *Bbc3 (*PUMA) / *Casp3, Ddit3 (*CHOP), *Eif2ak3, and Trib3*, well-known markers of ER stress. Transcripts downregulated in the CST-hCreb5 cluster were enriched for synaptic functions, consistent with prior work in other cell types linking axon injury to altered synaptic function^40–42^ (**Sup. Fig. 4**). Thus, after cervical axotomy a subset of CST neurons initiate a damage response that includes some activation of ER stress pathways and downregulation of synapse function. It should be emphasized that fewer than 20% of CST neurons displayed this response. In addition, this subset lacked other markers of cellular stress, notably *Atf3*, and did not upregulate regeneration-associated transcripts^7^ such as *Sox11, Tuba1a, Klf6, Gap43*, and others (**Supplemental Table 2**). Overall, although some signs of cellular stress can be detected in a subset of CST neurons, cervically injured CST neurons did not show widespread transcriptional changes or activation of axonal growth pathways.

### Intracortical injury, but not spinal injury, upregulates ATF3 and phospho-cJUN in CST neurons

DRG and RGC neurons, which display a much larger transcriptional response to axotomy than what we detected after spinal injury in CST neurons, both undergo a retrograde stress response that is marked by rapid phosphorylation of cJUN and upregulation of ATF3 protein^43^. We therefore compared these indicators in CST neurons at one and seven days after cervical axotomy to sham-injured controls. ATF3 was undetectable in any CST neuron at either time point (**Fig. 3c, d**) and phospho-cJUN was barely visible in most CST neurons (**Fig. 3e, f**). We noted, however, that at seven days post-injury some CST neurons showed faint phospho-cJUN signal. Quantification of signal intensity showed no significant difference between cervically injured and sham-injured controls (p>.05, one-way ANOVA with post-hoc Sidak’s), but a small population with elevated phospho-cJUN signal can be seen in the scatterplot at seven days (**Fig. 3f)**. In summary, in line with the snRNA-seq findings, detection of phospho-cJUN and ATF3 suggests a minimal cell body response to cervical axotomy in CST neurons.

**Figure 3:**
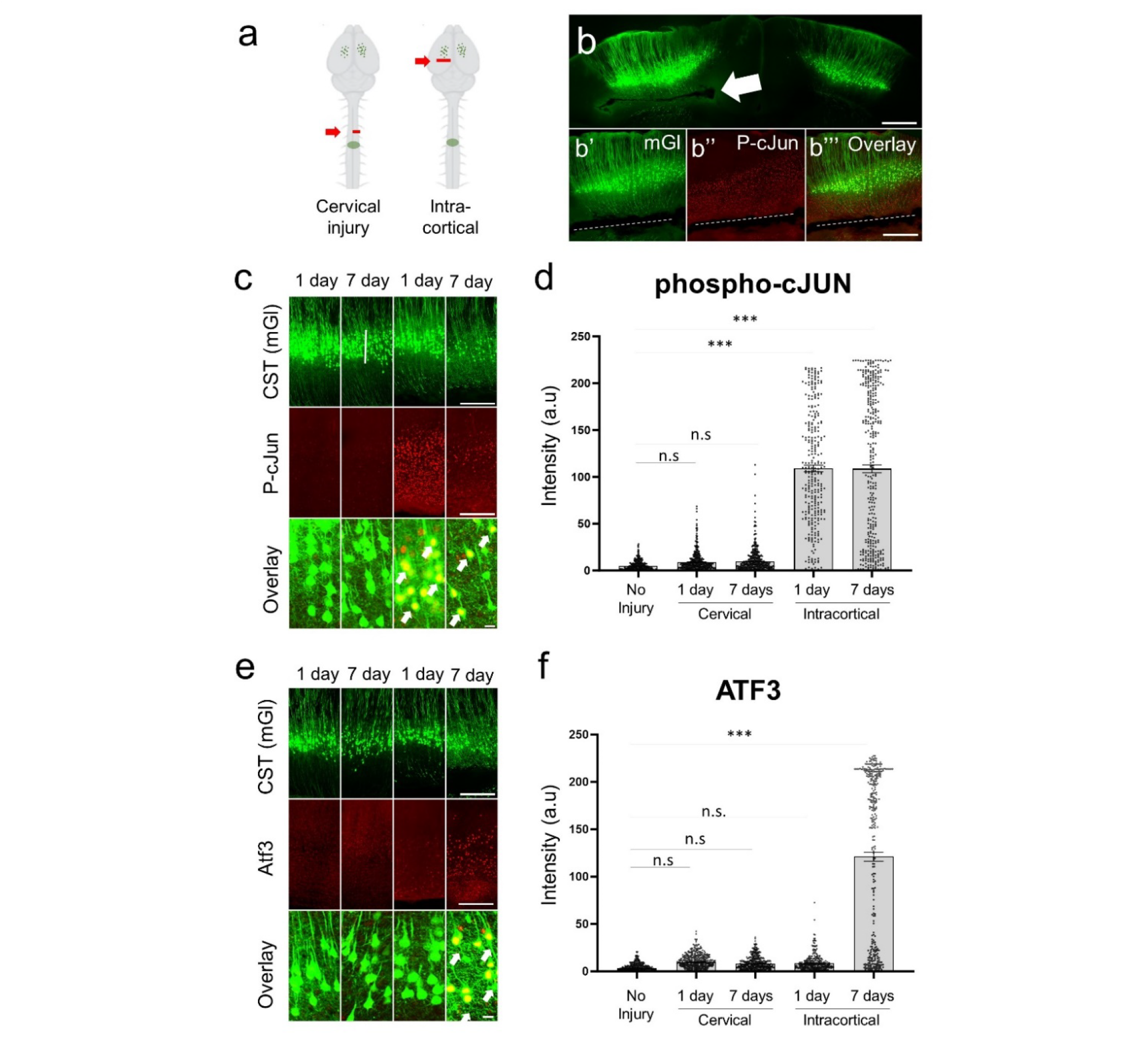
Intracortical but not spinal axon injury causes phosphorylation of cJUN and upregulation of ATF3 in CST neurons. (a) Experimental procedure. CST neurons were retrogradely labeled by Retro-AAV2-mGI and then injured in the cervical spinal cord or within the cortex, (b) shows a coronal section of cortex one week after intracortical injury with injury (arrow, dotted line) and nearby expression of p-cJUN, including in CST neurons (mGI, green), (c) Immunohistochemistry for p-cJUN one or seven days post injury, (d) Quantification of p-cJUN shows significant elevation after intracortical injury but not after spinal injury, (e) Immunohistochemistry for ATF3 (red) in CST neurons (green) one or seven days post injury, (f) Quantification of ATF3 signal shows significant elevation seven days after intracortical injury. Scale bars are 500pm (b). 200pm (upper panels, c, e), 10pm (lower panels c,e). ***p<.001, ANOVA with post-hoc Sidak’s.

Does this relatively subdued response reflect an inherent cellular difference in CST neurons, or does it arise from differences in the anatomical proximity to the site of injury? To distinguish these possibilities, we employed an intracortical injury model to sever descending CST axons within 1mm of their cell bodies and proximal to any collateral synapse formation (**Fig. 3a,3b, Sup. Fig. 5**). In striking contrast to the prior spinal injury, phospho-c-JUN was strongly upregulated at both one-and seven-days post-lesion, indicating rapid engagement of a retrograde stress response (**Fig 3c, d**). In addition, proximal injury resulted in strong expression of ATF3 by seven days post-lesion (**Fig. 3e, f**; p< .001, ANOVA with post-hoc Sidak’s). Importantly, all tissue processing and IHC procedures were performed simultaneously including both spinally and intracortically injured tissue, thereby strengthening the conclusion of a large difference in expression between the two conditions. We also noticed that by seven days after intracortical injury the number of retrogradely labeled CST neurons appeared diminished (compare 1-and 7-day panels in **Fig. 3c**). This visual impression was confirmed in a second cohort of animals in which CST neurons were imaged by tissue clearing and 3D imaging, showing a significant reduction in cell number in the weeks following intracortical injury (**Sup. Fig. 6)**. Thus, reminiscent of the RGC response to optic nerve injury and consistent with prior observations of CST cell loss after intracranial axotomy^44^, CST neurons appear to undergo significant cell death after proximal axotomy. Taken together, these data indicate that axotomies located near to CST cell bodies and proximal to collateral contacts with brain targets^45^ result in strong activation of cellular stress pathways and substantial cell death. These data rule out the possibility of cell-type insensitivity as the reason for the limited activation of phospho-c-Jun and Atf3 in CST neurons after spinal injury and instead indicate injury location as a determining factor.

### CST neurons show large transcriptional responses to intracortical injury

Because CST neurons upregulate damage-responsive phospho-cJUN and ATF3 after intracortical injury, we tested for a larger genome-wide response using snRNA-seq. As previously, CST cell nuclei were retrogradely labeled from the cervical spinal cord and seven days later a sharp blade was passed through subcortical white matter, severing CST axons approximately 1 mm from their cell bodies (**Fig. 4a, Supp. Fig. 5a**). At one, three, or seven days after injury Layer V cortex was micro-dissected under fluorescence, taking tissue only from brain sections in which visual inspection confirmed an injury located deep to layer V (**Supp. Fig. 5b**). In these experiments also employed DNA barcoding, delivered using AAV to CST neurons along with the nuclear reporter, which allowed pooling of injured and uninjured tissue and subsequent deconvolution. In UMAP plots, unlike the prior thoracic and cervical injury responses, proximally injured CST nuclei clustered separately from sham-injured nuclei and showed strong and near-universal upregulation of *Creb5* (**Fig. 4b**). Across the three time points more than 3000 DEGs significantly exceeded a two-fold threshold at either one, three, or seven days (**Supplemental Table 2**). Moreover, unlike in the prior spinal injury datasets these DEGs included many transcripts that were detected in a significant proportion of nuclei, indicating broad population-level participation in the transcriptional response (**Fig. 4c**). Network and gene ontology analyses of upregulated transcripts revealed enrichment of terms associated with wound healing, various inflammatory processes, adhesion, and actin cytoskeleton remodeling (**Fig. 4e,f**). Comparison of our data to a prior *in situ* hybridization-based characterization of CST gene expression after intracortical injury showed perfect correspondence across eight transcripts (*Atf3, Jun, Gap43, L1cam, Chl1, Stmn2* strongly upregulated in both, *Basp1 / Cap-23* and *Krox-24/Egr1* upregulated in neither)^33^. In summary, the large number of DEGs detected after intracortical injury confirms the sensitivity of the snRNA-seq approach in detecting transcriptional changes. This reaffirms the notion that the subdued response seen after spinal injury is indicative of a genuine biological signal rather than technical limitations.

**Figure 4:**
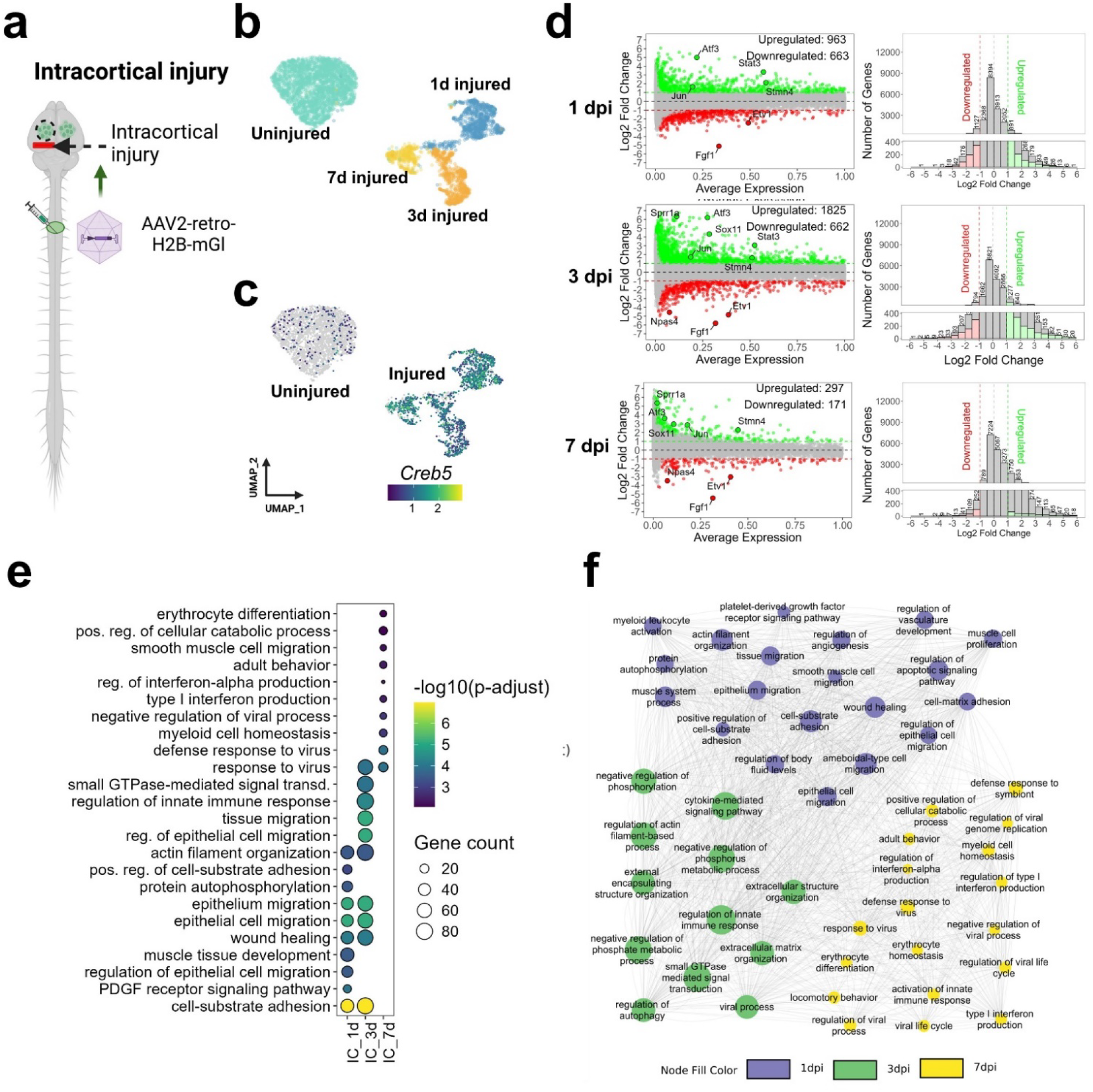
Single-nuclei profiling detects a large transcriptional response to intracortical injury in CST neurons. (A) Experimental design in which CST neurons were retrogradely labeled by AAV2-retro-H2B-mGI injection, axotomized in the lower cortex or left uninjured, and harvested for single-nuclei profiling 1, 3, or 7 days later, (b) UMAP clustering shows that injured samples at various post-injury intervals (1 day, 3 days, and 7 days) segregate from one another and from uninjured samples, (c) A feature plot showing elevated levels of *Creb*5 in intracortically injured CST neurons, (d) Histograms and MA plots show numerous transcripts significantly up-regulated (green) and down-regulated (red) in CST neurons after intracortical injury (p-value<0.05, non-parametric Wilcoxon rank sum test) (e) Gene ontology enrichment analysis indicating biological processes that are significantly over-represented in CST-upregulated gene sets at 1, 3, or 7 days post-injury (p-value<0.05, Benjamini-Hochberg test), (f) A gene ontology and pathway interaction network shows gene ontologies and pathways affected by intracorical injury, with the intensity of the node fill color representing the timeline (1 DPI (Violet), 3 DPI (Green), and 7 DPI (Yellow), p-value<0.05, Benjamini-Hochberg test))

Examination of stress-associated transcripts illustrates this phenomenon. CST-IC cells strongly upregulated ER-stress markers that were previously detected in the cervically injured CST-hCreb5 subset, notably *Bbc3 (*PUMA), *Casp3, Ddit3 (*CHOP), *Eif2ak3*, and *Trib3* (**Supplemental Table 2**). Besides these, intracortical injury caused strong upregulation of additional stress and apoptosis markers that were not affected by cervical injury, including *Atf3, Atf5, Pp1r15a, Gadd45g, Bcl2l11/BIM, Casp9*, and *Bcl6* (**Sup. Fig 7**). Thus, spinal injury appears to trigger only partial activation of the stress response;; we emphasize again that only a subset of ∼20% of cervically injured neurons display even this partial response. Intracortical injury also caused upregulation of well-known regeneration-associated genes, including transcription factors *Sox11, Stat3, Klf6, and Smad1*, and other RAGs such as *Gap43, L1cam, Sprr1a, Stmn2/SCG10, Gal, Inppk5*, and *Itga9* (**Sup. Fig. 7, Supplemental Table 2**). Of these, only *Itga9* was significantly upregulated after cervical spinal injury, and again only in the h*Creb5* subset.

The difference in scale raises the question of whether the responses are distinct at the molecular level or if they stem from a shared axotomy program that is activated less intensely by distant injury. The former model predicts minimal overlap between spinal and intracortical injury DEGs, at the level of chance, while the latter predicts DEGs that are shared by both injuries but with fold changes that are larger in the intracortical set. To distinguish these possibilities, we compared significantly regulated transcripts, using a relaxed 25% threshold (log2FC > 0.32) to sensitize detection. Of the 137 transcripts upregulated >25% in cervically injured CST neurons, 64% were also upregulated by intracortical injury, a significant enrichment (p<.001, hypergeometric test) (**Sup. Fig. 8a**). Overlap was even larger for the 317 transcripts upregulated in the hCreb5 group, with 78% upregulated by intracortical injury (p<.001, hypergeometric test). Fold change values in the common upregulated genes were positively correlated (R-squared = 0.4 and 0.6 for all CST and hCreb5 CST, respectively) with a slope significantly less than 1 (Δcervical / Δintracortical), indicating larger changes in the intracortical than spinal injury (**Sup. Fig. 8a**). Consistent with this, the intracortical fold change was larger than the spinal injury change in 97% and 85% of transcripts shared with the all-CST or hCreb subset, respectively. Overall, these data favor the existence of an axotomy response that is shared but which differs in the degree of activation, with spinal injury causing smaller changes in a subset of genes comprising of ∼10% of intracortical-responsive genes.

Finally, because these analyses relied on distinct sequencing libraries, we performed an additional experiment to ensure that variability in injury timing, library preparation, or sequencing were not driving the difference in DEG detection between SCI and IC samples. To enable pooling of different injury conditions in the same library, CST neurons in adult mice were retrogradely labeled by cervical injection of H2B-mGl combined with one of three distinct DNA barcodes. One week later the barcoded subgroups were subjected to cervical hemisection, intracortical injury, or remained uninjured, followed three days later by microdissection and flash freezing. Tissue from all three groups was then pooled such that nuclei isolation, FANS sorting, library preparation, and sequencing were identical. In the resulting data 94.2% of nuclei could be classified by barcode, with the remainders excluded for low or conflicting barcoding reads. The IC sample segregated from the SCI and uninjured nuclei in UMAP clustering (**Fig. 5a**) and in feature plots displayed strong elevation of stress and regeneration associated genes (RAGs) including *Creb5, Atf3, Sox11, Stat3, Klf6*, and *Sprr1a* compared to both uninjured and SCI samples (**Fig 5b**). Indeed, although SCI displayed only 33 DEGs compared to uninjured at a 2-fold threshold the IC sample had 1846 DEGs (**Fig. 5c,d; Supplemental Table 3**). Moreover, consistent with the prior IC datasets these DEGs included numerous additional RAGs including *Tubb2b, L1cam*, and *Gap43* (**Fig. 5d,e Supplemental Table 3**). In contrast, SCI nuclei did not exhibit significant upregulation of most RAGs. Thus, under identical conditions of FANS, library preparation, and sequencing, snRNA-seq detects a robust transcriptional response to IC injury but a muted response – approximately 50-fold lower – to SCI. Overall, these data show that when CST neurons experience a nearby injury stimulus they initiate a strong transcriptional response including upregulation of transcripts associated with axon regeneration. However, this response is largely absent following more distal axon injury.

**Figure 5:**
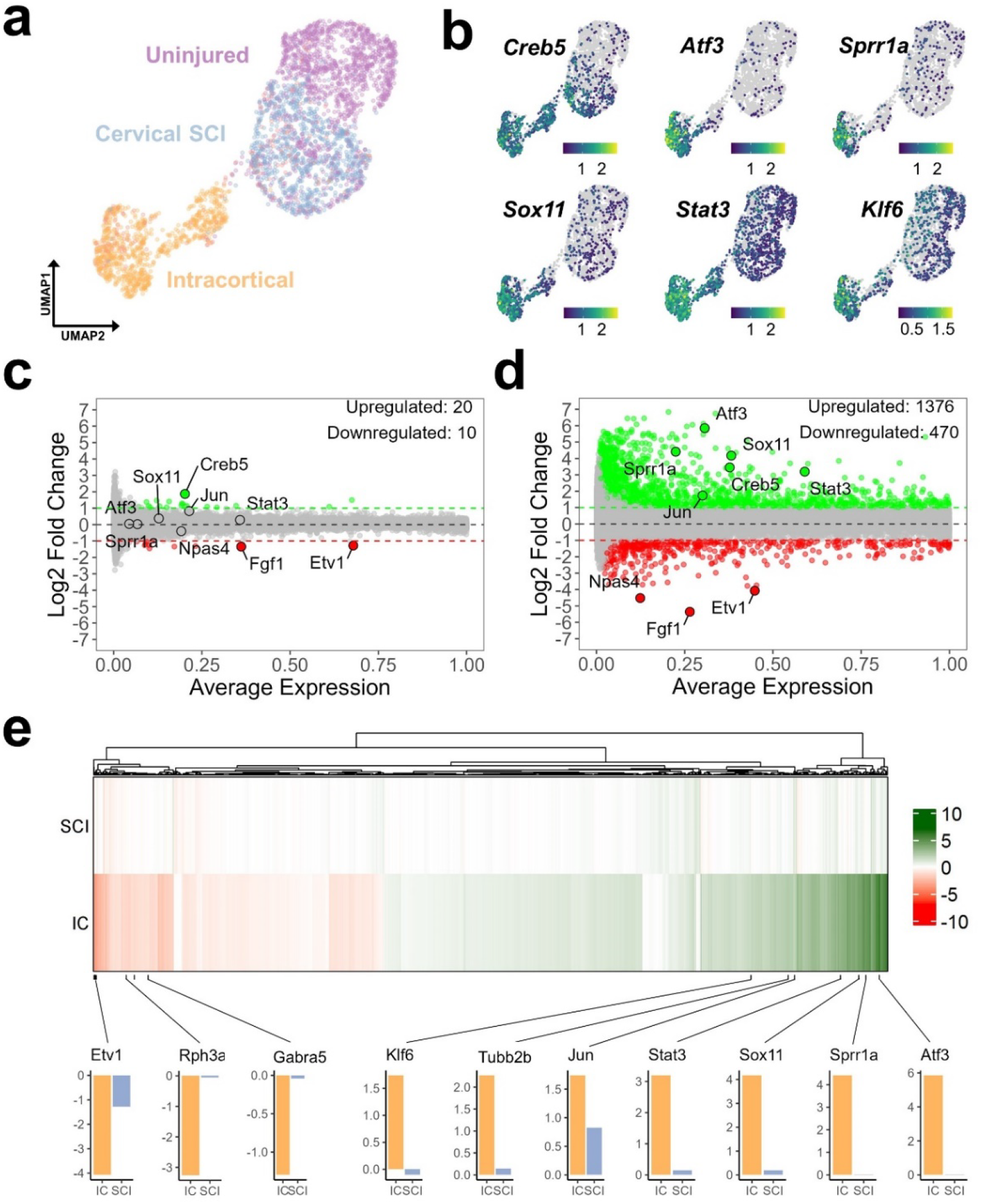
Matched within-library profiling confirms pronounced transcriptional changes in CST neurons after intracortical but not spinal injury. Animals received retrograde nuclear labeling along with a molecular barcode followed by no injury, spinal injury, or intracortical injury. Three days later tissue was pooled for library preparation, followed by post-sequencing deconvolution, (a) A UMAP plot shows the distribution of nuclei from uninjured (purple), cervical spinal cord injured (SCI, blue), and intracortical injury (orange) samples, (b) Feature plots illustrating strongly elevated expression of stress and regeneration-associated transcripts (*Creb5, Atf3, Sprrla, Soxll, Stat3, Klf6*) in CST neurons after intracortical injury. (c,d) MA plots show fewer upregulated (green) and downregulated (red) in CST neurons that received spinal injury (C) compared to intracortical injury (d) (p-value<0.05, non-parametric Wilcoxon rank sum test) (e) A heatmap compares the log fold change in gene expression between spinal and intracortical injury, with a scale from green (upregulation) to red (downregulation). Bottom, bar graphs comparing the log fold change of specific growth-associated genes between intracortical (IC) and cervical SCI conditions.

### Proximally injured CST neurons share gene changes with injured RGC and DRG neurons

We next examined the extent to which the response of CST neurons to nearby axotomy resembles that of RGC or DRG neurons. To match our data, we assembled sets of transcripts that are significantly up-and down-regulated at a two-fold threshold within the first week after axon injury from optic nerve crush ^11^(RGC) or spinal axotomy (DRG)^10^. Intersection of the datasets showed that of 2285 transcripts upregulated by CST neurons between 1-and 7-days post-injury, 569 and 329 were shared with axotomy-upregulated transcripts in DRG or RGC neurons, respectively, significantly more than chance (p < 3.495e-106 and p < 2.670e-92, hypergeometric test**) (Fig. 6a, Supplemental Table 4)**. Notably, despite the differences in cell types, injuries, and experimental techniques, 145 transcripts were upregulated in all three datasets, more than ten times the number predicted by chance. Thus, although many axotomy-upregulated transcripts are unique to the cell type and injury, there also exists a shared set of transcripts that represent a common response to injury.

**Figure 6:**
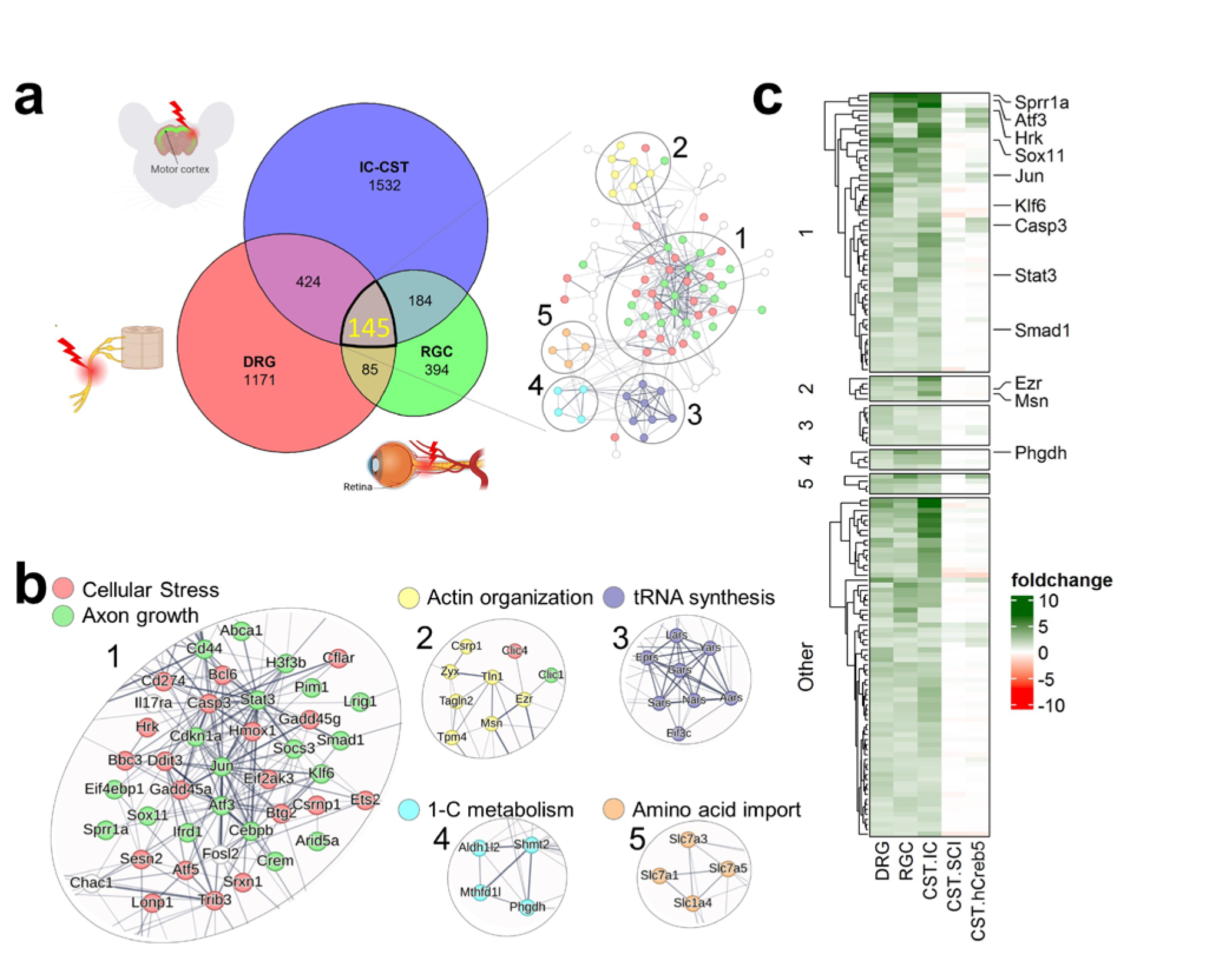
Key growth-related genes are activated in response to injuries across various cell types. (a) A Venn diagram shows the overlap of differentially expressed genes In CST neurons after intracortical injury (IC), dorsal root ganglia (DRG) after spinal injury, and retinal ganglion cells (RGC) after optic nerve crush, indicating unique and shared transcriptional responses, (b) Interaction network (String) networks of the genes shared among the three injury sub-types reveal five distinct functional categories: (1) Cellular Stress and Axon Growth, (2) Actin Interaction, (3) tRNA Synthesis, (4) 1-C metabolism, and (5) Amino Acid Import. Each network node represents a gene, with edges indicating interactions or functional associations of all the 145 genes found in the all 3 groups.(c) A heatmap displaying the gene expression changes across different pathways in the DRG, RGC, and CST after intracortical (IC) or spinal injury (SCI). The color scale reflects the fold change from downregulation (red) to upregulation (green). Specific genes of interest are annotated on the side, highlighting their expression patterns across the conditions.

For insight into the function of the 145 commonly upregulated (CU) transcripts we generated a functional interaction network using STRING^46^ (**Fig. 6a,b**). Five main clusters emerged, four of which contained between four and eight gene products based on actin organization, tRNA synthesis, amino acid import, and 1-carbon metabolism. The fifth and largest cluster contained numerous RAGs including *Sox11, Smad1, Klf6, Stat3, Cdkn1a*, and *Sprr1a* and stress-related transcripts such as *Ddit3 (Chop), Hrk, Eifak3 (Perk), and Bbc3* (**Fig. 6b**). To systematically identify transcripts that could contribute functionally to axon growth, each of the 145 CU gene symbols was combined with “axon” in PubMed and the resulting manuscripts manually examined for evidence that knockout or overexpression in any neuronal cell type resulted in a change in axon length (**Supplemental Table 5**). Interestingly, 32 of 145 (22%) of transcripts met these criteria, with all but five playing a positive role in axon growth (evidence for increased growth upon overexpression and/or decreased growth after knockdown). Similarly, all gene symbols were combined with “neuron” and “apoptosis” and the results manually screened for experimental evidence of neuron-intrinsic effects on cell survival, yielding 40 (27.6%) with a functional role in cell survival and/or integrated stress responses. For comparison, in a background set of 145 transcripts upregulated in the CST-IC set but not shared by DRG or RGC neurons only 11 could be manually linked to axon growth or neuronal survival, significantly lower than detection in the CU set (p<.0001, Chi-squared) (**Supplemental Table 5**). Finally, we examined the behavior of CU transcripts in the prior spinal injury sets. Across all CST neurons after cervical injury, only one CU transcript, *Hrk*, was significantly upregulated more than 2-fold, with nine additional transcripts showing upregulation that was statistically significant but below this fold threshold (**Fig. 6c**). Within the hCreb5 subset, 14 CU transcripts were upregulated more than two-fold, including four axon-linked terms (*Jun, Arid5a, Rhoq, Abca1*) and five stress-related terms (*Hrk, Trib3, Bbc3, Eif2ak3, Casp3*, and *Ddit3*) (**Supplemental Table 4, Fig. 6c**). Besides these, nine CU transcripts showed an injury-triggered increase that was significant but below a two-fold threshold. It is notable, however, that 122 (84%) of CU transcripts were unaffected by spinal injury even in this higher-responding subset. Overall, these data identify a common set of axon-and stress-relevant transcripts that are strongly upregulated by CST neurons after proximal axon injury but minimally after spinal injury.

## Discussion

We have analyzed the transcriptional responses of mixed supraspinal populations to thoracic spinal injury and the response of CST neurons to thoracic, cervical, and intracortical injury. The main finding is that spinal injury led to minor transcriptional changes, as evidenced by the number of affected transcripts and the degree of up-or down-regulation in affected transcripts. Conversely, axotomy located close to CST cell bodies resulted in significantly greater transcriptional effects. These changes were comparable to those observed in other cell types, suggesting that limitations in cell-type or technical sensitivity are not the primary explanations for the low response to spinal injury. Instead, it is more likely that the cellular response to spinal injury is limited by anatomical factors, such as the intervening distance to the injury site or the preservation of proximal collateral connections. Furthermore, when comparing the CST response to proximal injury with that of other cell types, we identified a central network of injury response involving many transcripts linked with cellular stress and axon growth. However, this network is only minimally activated by spinal injury. Thus, the limited regenerative response of descending neurons to spinal injury is more likely caused by the failure to detect or react to distant injury, rather than inherent cellular constraints or an inability to maintain a transient reversion to an embryonic state.

This conclusion correlates closely with findings from decades past, emphasizing the critical role of injury proximity in the process of axon regeneration. Seminal work in the 1980s established that although some supraspinal populations can regenerate axons into peripheral nerve grafts placed into cervical spinal cord, this ability was mostly lost when grafts were placed in thoracic regions ^47^. Similarly, the ability of RGCs to regenerate axons into peripheral nerve grafts requires that injury occur within about three millimeters of the cell body ^48^. Likewise, a series of manuscripts in the 1990s showed that in rubrospinal neurons both regenerative ability and the expression of regeneration-associated genes depends on injury proximity ^32,49,50^. Finally, research in CST neurons identified six RAGs that increase in expression after intracortical but not spinal injury; our work not only replicates this result but also extends it across the genome^33^. Therefore, our findings of relatively modest transcriptional responses to spinal injury reinforce and expand upon an earlier concept that the proximity of injury plays an important role in regulating the regenerative response of neurons.

Our conclusions, however, are unexpected considering a recent study examining post-injury transcriptional patterns in CST neurons^9^. This study reported approximately ten times more DEGs at similar fold change thresholds after cervical spinal cord injury, suggesting a spontaneous, albeit transient, reversion to an embryonic pattern of gene expression after spinal injury^9^. Technical differences between the studies might contribute to the discrepancy, such as the ten-day post-injury timepoint in the previous study compared to seven days in our study and, crucially, the use of a TRAP approach^51^ versus our single-nuclei analysis centered approach. The TRAP approach could potentially detect injury-induced changes in RNA translation not reflected in altered mRNA levels within the nuclear compartment. Interestingly, one area of agreement between the two datasets concerns the injury-responsive transcripts that we found to be shared across cell types (**Fig. 5**). Of the 145 transcripts upregulated two-fold in intracortical-CST, DRG, and RGC datasets, Poplawski *et al*. found only 12 (8%) to be upregulated after cervical injury, none of which have previously been linked to axon growth. Thus, both our data and those from Poplawski *et al*. converge on the critical point that after spinal injury, CST neurons fail to strongly upregulate a core set of pro-regenerative transcripts (e.g. *Atf3, Sox11, Klf6, Smad1, Stat3, Gap43, L1cam, Gal*, etc).

Indeed, the lack of ATF3 expression and the very low level of c-JUN phosphorylation observed in CST neurons after spinal injury, as confirmed at the protein level through Immunohistochemistry (IHC), may play a significant role in understanding the low regenerative response. ATF3 plays a crucial role in regulating axon regeneration, as evidenced by experiments across multiple cell types showing that genetic ablation of *Atf3* substantially reduces regenerative axon growth^52 28,31–33^. More broadly, JUN phosphorylation and ATF3 upregulation are markers of damage-sensing pathways, notably the DLK pathway, which is critical for initiating axon growth in various cell types^2,53,54^. Studies conducted across a wide range of species and cell types indicate that blocking the DLK pathway essentially abolishes injury-triggered axon growth, indicating that this pathway may be almost universally required for neurons to sense and respond to injury^2,55,56^. In this context, our findings that DLK indicator genes are either low or absent in CST neurons following spinal injury hint that a muted activation of damage-sensing pathways may limit their potential for a regenerative response.

In summary, our thorough analysis of transcriptional responses in mixed supraspinal populations and CST neurons following different spinal injuries reveals modest changes in gene expression, particularly in response to distal spinal injuries. This redirects attention to an earlier hypothesis that anatomical factors, particularly the proximity of the injury to the neuron’s cell body, play a crucial role in determining the extent of transcriptional response and regeneration. Additionally, our data confirm that CST neurons do not spontaneously regress to a growth-competent state following spinal injury, underscoring the need to therapeutically stimulate a transcriptional program supportive of growth^6,57– 59^. Furthermore, the minimal activation of essential damage-sensing pathways, such as the DLK pathway—as indicated by the sparse presence of DLK pathway genes—suggests a potential mechanistic explanation for the restrained regenerative responses observed in CST neurons after spinal injuries. Together, these insights enhance our understanding of the regulatory mechanisms influencing axon regeneration and pave the way for further research into the biological processes constraining neuronal recovery following injury.

## Methods

### Mice and husbandry conditions

All procedures involving animals were conducted in strict adherence to ethical guidelines provided by the Guide for the Care and Use of Laboratory Animals published by the National Institutes of Health (NIH). The Institutional Animal Care and Use Committee (IACUC) at Marquette University reviewed and approved all animal experimental protocols (approval numbers 3283, 4013). Housing conditions for the mice included a controlled 24-hour cycle of 12 hours of light followed by 12 hours of darkness, temperature of 22°C ± 2°C, and relative humidity between 40% and 60%.

### AAV preparation

rAAV2-retro-CAG-H2B-mGreenLantern (Addgene#177332) was produced at the University of Miami viral core facility at the Miami Project to Cure Paralysis, titer = 1.4×10^13^ particles/ml. rAAV2-retro-H2B-mScarlet (Addgene#191093) and rAAV2-retro-tdTomato (Addgene #59462) were produced at the University of North Carolina Viral Vector Core at 6×10^12^ particles/ml and 3×10^12^ particles/ml respectively. To create barcoding constructs a 625bp sequence corresponding to positions 6903-7525 of human Malat RNA, previously identified as responsible for nuclear localization, was synthesized by Genscript with a 3’ BamHI and a 5’ SalI-barcode-NotI cassettes (designated Malat-BC). Using rAAV2-H2B-mGl (above), BamHI and SalI were used to replace the mGl-WPRE region with Malat-BC. Barcode sequences of 20nt (BC0 and BC1) or 100nt (BC102) were randomly generated and sequenced with flanking SalI and NotI sites, which were used for insertion into the CAG-Malat-BC, generating CAG-Malat-BC0, -BC1, and BC102 and used for AAV production at the Univ. of Miami Viral Vector Core. All viruses were diluted to 2.5×10^12^ with PBS immediately prior to injection.

### Surgical procedures

All surgeries were performed under ketamine/xylazine anesthesia using adult C57/Bl6 mice 8-12 weeks of age (20-28g). For AAV injections mice were mounted on a custom spine stabilizer and laminectomy performed at C5 or T10-12. AAV-retro particles (1µl, 2.5×10^12^ particles/ml) were injected by a 1701 Hamilton syringe fitted with a pulled glass capillary needle and driven by a Stoelting QSI pump (catalog #53311), guided by a micromanipulator. AAV was injected at a rate of 0.04 μl/min, 0.35 mm lateral to the midline at an initial depth of 0.8 (500µl) and then raised to 0.6 mm (additional 500µl).

Thoracic crush injuries were performed as in Wang et al. 2022. Mice were mounted on a custom spine stabilization device, laminectomy was performed at thoracic vertebra 10-12, and forceps with a stopper of 0.15mm width was used to compress the spinal cord for 15 seconds, flipped in orientation, and reapplied at the same site for an additional 15 seconds. For cervical transection, as in our previous work^23^, mice were mounted in a custom spine stabilizer and a vibraknife device was used to produce a transection that extended from slightly left of the midline to the right lateral edge of the spinal cord and to a depth of 0.85mm. This injury severs both dorsal and dorsal-lateral populations of CST axons. Finally, intracortical injuries were performed with procedures modified from previously published work^33^. Mice were mounted on a stereotaxic frame in earbars, and the left cortex exposed by gentle scraping of the skull with #11 scalpel blade. A 30G needle was bent at 90° 2mm from the tip and with stereotactic guidance was placed on the surface of the cortex with the bent tip in the sagittal plane, 0.5mm to the left of the midline, and with the tip aimed caudally. The needle was then lowered 0.6mm into the brain and rotated 180° clockwise, swinging the tip of the needle beneath left motor cortex, and then rotated back to the starting position and withdrawn.

### Nuclei isolation, library preparation, and sequencing

Following previously published procedures^34^ mice were isoflurane-anesthetized, decapitated, and brains rapidly placed in ice-cold slushy artificial cerebrospinal fluid (Hearing et al., 2013) for one minute. Working quickly on ice, brains were then sectioned in the sagittal (thoracic injury) or coronal (cervical or intracortical injury) plane at 500-micron intervals using Adult Mouse Brain Slicer Matrix on ice (Zivic Instruments BSMAS005-2). Retrogradely labeled populations were rapidly microdissected in ACSF using a stereomicroscope and fluorescence adapter (NIGHTSEA SFA-GR) and immediately frozen on dry ice; collected target regions included corticospinal for all treatments and also hypothalamic, midbrain, pontine, and medullary supraspinal populations for the thoracic injury group. Samples were then stored at -80 °C for up to one month until FANS sorting.

Cell nuclei were isolated and sorted on the day of library preparation as previously described ^34,60^. Briefly, tissue was dounced in Nuclei EZ Lysis Buffer (Sigma-Aldrich N 3408), incubated on ice for five minutes, centrifuged at 500 G at 4 °C for five minutes, resuspended 4ml Nuclei EZ Lysis Buffer, incubated on ice, and centrifuged at 500G at 4 °C for 5 minutes. The resulting pellet was resuspended in 500uL of Nuclei Suspension Buffer (2% BSA, 40U/uL RNase Inhibitor (Invitrogen Ref. AM2684), 1X PBS), passed through a 20um filter and moved directly to the FACS machine. The collection tube for the sorted nuclei was coated with 5% BSA and contained the 10X Genomics master mix. Debris and doublets were gated out using side scatter area (SSC-A), side scatter width (SSC-W), forward scatter area (FSC-A) and forward scatter width (FSC-W) and then collected based on level of fluorescence so that only the brightest were collected. With the above parameters, the FACS machine was set to collect 5,000 nuclei in five to seven minutes. The collected nuclei were then prepared into libraries, using Chromium Next GEM Single Cell 3’ Reagent Kits v3.1 (PN-1000269), according to the manufacture protocol (10X Genomics, CG000204 Rev D).

Sequencing was performed at the UW-Madison Biotechnology Center using an Illumina NovaSeq 6000, yielding at least 400 reads per library. Files were processed with CellRanger using default parameters to produce a Unique Molecular Identifier (UMI) matrix for all nuclei-containing droplets. Normalization, dimensionality reduction, and cell clustering were performed using the Seurat package version 5.0. (https://www.nature.com/articles/s41467-022-35574-x#ref-CR18)^61^. Cell clusters were identified by performing principal component analysis (PCA) on the top 30 principal components, with significant components determined by an elbow plot and jackstraw test. Cells were clustered using the Louvain method in the FindClusters function (resolution = 0.05) and visualized using Uniform Manifold Approximation and Projection (UMAP). For injured versus uninjured data comparison, the “merge” and “FindIntegrationAnchors” functions from SEURAT were used to integrate the datasets. Differential expression analysis was conducted using the “FindMarkers” function in SEURAT, applying a non-parametric Wilcoxon rank-sum test. Cluster plots in Seurat were generated using the “scCustomize” package in R (https://github.com/samuel-marsh/scCustomize), while MA-plots, histograms, and heatmaps were created using the “ggplot2” package (https://ggplot2.tidyverse.org/)^62^. we have utilized ShinyCell^63^ in R to provide prebuilt clusters of the analyzed scRNA-seq data, enabling direct visualization and interactive exploration. The clustered data is publicly accessible and can be explored through our dedicated webpage at https://venkateshlab.github.io/CNS_response/.

### Gene Ontology and Network Analysis

Gene ontology analysis was conducted using the “ClusterProfiler” (https://www.ncbi.nlm.nih.gov/pmc/articles/PMC3339379/) package in R ^64,65^, with input genes showing a fold change value ≥1 or ≤-1 and an adjusted p-value <0.05, marking them as upregulated and downregulated, respectively. The “org.Mm.eg.db” database served as the reference for ontology searches. The analysis included Biological Process (BP), Cellular Component (CC), and Molecular Function (MF) categories, setting gene count size limits between 3 and 500, and a p-value cutoff of <0.05. The top 20 pathways were visualized using the dot plot function from the enrich plot package in R, and the results were saved as a CSV file for network visualization in Cytoscape (3.10)^66^ (https://www.ncbi.nlm.nih.gov/pmc/articles/PMC403769/).

### Tissue clearing and imaging

Three weeks after cervical AAV injection and one to fourteen days after sham or intracortical injury animals underwent transcardial perfusion with 0.9% saline and 4% paraformaldehyde (PFA) solutions in 1× phosphate-buffered saline (PBS) (15710, Electron Microscopy Sciences, Hatfield, PA). Whole brains were dissected and fixed overnight in 4% PFA at 4°C and washed three times in PBS pH 7.4, in PBS. and the brains were cleared using a modified version of the 3DISCO ^67,68^. Whole mouse brains were incubated on a rotating shaker at room temperature in 50%, 80%, and twice with 100% peroxide-free tetrahydrofuran (THF; Sigma-Aldrich, 401757) for 12 hr each for a total of 2 days. Samples were transferred to BABB solution (1:2 ratio of benzyl alcohol, Sigma-Aldrich, 305197; and benzyl benzoate, Sigma-Aldrich, B6630) for three hours. After clearing, whole brains were imaged the same using a high-speed confocal, Andor Dragonfly 202-2540 (Oxford Instruments), based on a Leica microscope. We used a 10× Plan Apo, NA 0.45 and W.D 2.8. The laser was a solid-state 488 nm diode laser at 150 mW. Laser power was set at 15% and exposure time for each plane was between 60 ms. The camera used was a Sona sCMOS 4.2B-6 set at ROI size (W × H) at 2048 × 2048. All images were stitched using Imaris Stitcher Vx64 10 (Oxford Instruments). After stitching, 3D rendering and spot detection were performed using Imaris 10.01. The number of spots detected in cortical tissue dorsal to the region of intracortical injury was quantified and then normalized to an identically sized region in the contralateral (uninjured) cortex.

### Tissue processing and immunohistochemistry

For assessment of spinal injuries, at the time of brain microdissection the spinal cord was rapidly dissected and immersed in 4% paraformaldehyde (PFA) in 1× phosphate-buffered saline (PBS) (15710-Electron Microscopy Sciences, Hatfield, PA). For assessment of phospho-cJUN and ATF3 expression animals were perfused with 4% PFA, after which brains and spinal cords removed and post-fixed overnight in 4% PFA at 4°C. Spinal cords and cortical tissues were embedded in 12% gelatin in 1× PBS (G2500-Sigma Aldrich, St.Louis, MO) and cut via Vibratome to yield 100 μm sections. Sections were incubated overnight with primary antibodies GFAP (DAKO, Z0334 1 : 500, RRID:AB_10013482), p-cJUN (Cell Signaling 9261 1:500, RRID:AB_2130162, or ATF3 (Atlas Antibodies Cat#HPA001562, RRID:AB_1078233) and rinsed and then incubated for 2 h with appropriate Alexa Fluor-conjugated secondary antibodies (R37117, Thermofisher, Waltham, MA, 1 : 500). Fluorescent images were acquired using a Nikon A1R+ laser scanning confocal microscopy system on a Nikon Ti2-E inverted microscope. For quantification of cJUN and ATF3, regions of interest (ROIs) were first manually drawn in NIS Elements software using the mScarlet label within retrogradely transduced CST cell bodies. The average pixel intensity of the detection channel was then determined for each ROI. All visible CST cell bodies from at least two replicate cortical sections for each animal were selected by observers blinded to treatment.

## Acknowledgments

This work was supported by grants from NINDS R01 NS083983, The Bryon Riesch Paralysis Foundation, the Christopher and Dana Reeve Foundation (Awarded to Murray Blackmore) and funds from CSIR-CCMB, SERB-SRG and CSIR (Awarded to Ishwariya Venkatesh)

## Data availability

There are no restrictions on data availability. All the datasets generated and analyzed during the current study are available at figshare https://doi.org/10.6084/m9.figshare.c.7238734.v1 The clustered data is publicly accessible and can be explored through our dedicated webpage at https://venkateshlab.github.io/CNS_response/.

## Code availability

There was minimal custom code development and all software used in data analyses are previously published, open access, and have been cited under the relevant methods section. Links to relevant software repositories/documentation is listed here.

Cytoscape^66^ - https://cytoscape.org/

Cellranger^69^ - https://github.com/10XGenomics/cellranger.

SEURAT^61^ - https://github.com/satijalab/seurat;

clusterProfiler^64,65^ - https://bioconductor.org/packages/release/bioc/html/clusterProfiler.html;

ggplot^62^ - https://ggplot2.tidyverse.org/

scCustomize - https://github.com/samuel-marsh/scCustomize

Manual plots - https://github.com/RegenerationLab/Supraspinal_Injury

scRNA cluster - https://venkateshlab.github.io/CNS_response/

ShinyCell^63^ - https://github.com/SGDDNB/ShinyCell

## Notes

**Conflict of Interest:** None

https://doi.org/10.6084/m9.figshare.c.7238734.v1

https://github.com/RegenerationLab/Supraspinal_Injury

https://venkateshlab.github.io/CNS_response/

### Competing Interest Statement

The authors have declared no competing interest.

### Summary of Updates

This version of the manuscript has been revised to correct small errors in the text, to include links to datasets, and to update supplementary data.

## References

1. Bareyre, F. M. et al. The injured spinal cord spontaneously forms a new intraspinal circuit in adult rats. Nat Neurosci 7, 269–277 (2004).

2. Watkins, T. A. et al. DLK initiates a transcriptional program that couples apoptotic and regenerative responses to axonal injury. Proc Natl Acad Sci U S A 110, 4039–4044 (2013).

3. Hassannejad, Z. et al. The fate of neurons after traumatic spinal cord injury in rats: A systematic review. Iran J Basic Med Sci 21, 546–557 (2018).

4. Varadarajan, S. G., Hunyara, J. L., Hamilton, N. R., Kolodkin, A. L. & Huberman, A. D. Central nervous system regeneration. Cell 185, 77–94 (2022).

5. He, Z. & Jin, Y. Intrinsic Control of Axon Regeneration. Neuron 90, 437–451 (2016).

6. Venkatesh, I. & Blackmore, M. G. Selecting optimal combinations of transcription factors to promote axon regeneration: Why mechanisms matter. Neuroscience Letters vol. 652 64–73 Preprint at 10.1016/j.neulet.2016.12.032 (2017).

7. Ma, T. C. & Willis, D. E. What makes a RAG regeneration associated? Front Mol Neurosci 8, 43 (2015).

8. Zhang, Y., Zhao, Q., Chen, Q., Xu, L. & Yi, S. Transcriptional Control of Peripheral Nerve Regeneration. Mol Neurobiol 60, 329–341 (2023).

9. Poplawski, G. H. D. et al. Injured adult neurons regress to an embryonic transcriptional growth state. Nature 2020 581:7806 581, 77–82 (2020).

10. Renthal, W. et al. Transcriptional Reprogramming of Distinct Peripheral Sensory Neuron Subtypes after Axonal Injury. Neuron 108, 128-144.e9 (2020).

11. Tran, N. M. et al. Single-Cell Profiles of Retinal Ganglion Cells Differing in Resilience to Injury Reveal Neuroprotective Genes. Neuron 104, 1039-1055.e12 (2019).

12. Dhara, S. P. et al. Cellular reprogramming for successful CNS axon regeneration is driven by a temporally changing cast of transcription factors. Scientific Reports 2019 9:1 9, 1–12 (2019).

13. Matson, K. J. E. et al. Single cell atlas of spinal cord injury in mice reveals a pro-regenerative signature in spinocerebellar neurons. Nat Commun 13, (2022).

14. Fink, K. L., López-Giráldez, F., Kim, I.-J., Strittmatter, S. M. & Cafferty, W. B. J. Identification of Intrinsic Axon Growth Modulators for Intact CNS Neurons after Injury. Cell Rep 18, 2687–2701 (2017).

15. Li, S. et al. The transcriptional landscape of dorsal root ganglia after sciatic nerve transection. Sci Rep 5, 16888 (2015).

16. Xu, L. et al. Integrated analyses reveal evolutionarily conserved and specific injury response genes in dorsal root ganglion. Sci Data 9, (2022).

17. Palmisano, I. et al. Epigenomic signatures underpin the axonal regenerative ability of dorsal root ganglia sensory neurons. Nat Neurosci 22, 1913–1924 (2019).

18. Tian, F. et al. Core transcription programs controlling injury-induced neurodegeneration of retinal ganglion cells. Neuron 110, 2607-2624.e8 (2022).

19. Jacobi, A. et al. Overlapping transcriptional programs promote survival and axonal regeneration of injured retinal ganglion cells. Neuron 110, 2625-2645.e7 (2022).

20. Park, K. K. et al. Promoting axon regeneration in the adult CNS by modulation of the PTEN/mTOR pathway. Science 322, 963–6 (2008).

21. Sun, F. et al. Sustained axon regeneration induced by co-deletion of PTEN and SOCS3. Nature 480, 372–375 (2011).

22. Belin, S. et al. Injury-Induced Decline of Intrinsic Regenerative Ability Revealed by Quantitative Proteomics. Neuron 86, 1000–1014 (2015).

23. Norsworthy, M. W. et al. Sox11 Expression Promotes Regeneration of Some Retinal Ganglion Cell Types but Kills Others. Neuron 94, 1112-1120.e4 (2017).

24. Galvao, J. et al. The Krüppel-Like Factor Gene Target Dusp14 Regulates Axon Growth and Regeneration. Investigative Opthalmology & Visual Science 59, 2736 (2018).

25. Feng, Q., Wong, K. A. & Benowitz, L. I. Full-length optic nerve regeneration in the absence of genetic manipulations. JCI Insight (2023) doi:10.1172/JCI.INSIGHT.164579.

26. Fernandes, K. A., Harder, J. M., Kim, J. & Libby, R. T. JUN regulates early transcriptional responses to axonal injury in retinal ganglion cells. Exp Eye Res 112, 106–117 (2013).

27. Fawcett, J. W. The Struggle to Make CNS Axons Regenerate: Why Has It Been so Difficult? Neurochem Res 45, 144 (2020).

28. Starkey, M. L. et al. Assessing behavioural function following a pyramidotomy lesion of the corticospinal tract in adult mice. Exp Neurol 195, 524–539 (2005).

29. Moreno-López, Y., Olivares-Moreno, R., Cordero-Erausquin, M. & Rojas-Piloni, G. Sensorimotor Integration by Corticospinal System. Front Neuroanat 10, 24 (2016).

30. Lemon, R. N. Descending Pathways in Motor Control. Annu Rev Neurosci 31, 195–218 (2008).

31. Blackmore, M., Batsel, E. & Tsoulfas, P. Widening spinal injury research to consider all supraspinal cell types: Why we must and how we can. Exp Neurol 346, 113862 (2021).

32. Fernandes, K. J. L., Fan, D.-P., Tsui, B. J., Cassar, S. L. & Tetzlaff, W. Influence of the Axotomy to Cell Body Distance in Rat Rubrospinal and Spinal Motoneurons: Differential Regulation of GAP-43, Tubulins, and Neurofilament-M. J. Comp. Neurol 414, 495–510 (1999).

33. Mason, M. R. J., Lieberman, A. R. & Anderson, P. N. Corticospinal neurons up-regulate a range of growth-associated genes following intracortical, but not spinal, axotomy. Eur J Neurosci 18, 789–802 (2003).

34. Beine, Z., Wang, Z., Tsoulfas, P. & Blackmore, M. G. Single nuclei analyses reveal transcriptional profiles and marker genes for diverse supraspinal populations. J Neurosci JN-RM-1197-22 (2022) doi:10.1523/JNEUROSCI.1197-22.2022.

35. Wang, Z. et al. Brain-wide analysis of the supraspinal connectome reveals anatomical correlates to functional recovery after spinal injury. Elife 11, (2022).

36. Fink, K. L., Strittmatter, S. M. & Cafferty, W. B. J. Comprehensive Corticospinal Labeling with mu-crystallin Transgene Reveals Axon Regeneration after Spinal Cord Trauma in ngr1-/-Mice. J Neurosci 35, 15403–18 (2015).

37. Yao, S. Q., Wang, M., Liang, J. J., Ng, T. K. & Cen, L. P. Retinal transcriptome of neonatal mice after optic nerve injury. PLoS One 18, (2023).

38. DeVault, L. et al. The response of Dual-leucine zipper kinase (DLK) to nocodazole: Evidence for a homeostatic cytoskeletal repair mechanism. PLoS One 19, (2024).

39. Zhang, M. et al. Spatially resolved cell atlas of the mouse primary motor cortex by MERFISH. Nature 2021 598:7879 598, 137–143 (2021).

40. Hilton, B. J. et al. An active vesicle priming machinery suppresses axon regeneration upon adult CNS injury. Neuron 110, 51 (2022).

41. Tedeschi, A. et al. The Calcium Channel Subunit Alpha2delta2 Suppresses Axon Regeneration in the Adult CNS. Neuron 92, 419–434 (2016).

42. Kiyoshi, C. & Tedeschi, A. Axon growth and synaptic function: A balancing act for axonal regeneration and neuronal circuit formation in CNS trauma and disease. Dev Neurobiol 80, 277–301 (2020).

43. Tsujino, H. et al. Activating transcription factor 3 (ATF3) induction by axotomy in sensory and motoneurons: A novel neuronal marker of nerve injury. Mol Cell Neurosci 15, 170–182 (2000).

44. Giehl’, K. M. & Tetzlaff2, W. BDNF and NT-3, but not NGF, Prevent Axotomy-induced Death of Rat Corticospinal Neurons In Vivo. European Journal of Neuroscience 8, 1167–1175 (1996).

45. Sinopoulou, E. et al. Rhesus macaque versus rat divergence in the corticospinal projectome. Neuron 110, 2970-2983.e4 (2022).

46. Szklarczyk, D. et al. STRING v10: protein-protein interaction networks, integrated over the tree of life. Nucleic Acids Res 43, D447–D452 (2015).

47. Richardson, P. M., Issa, V. M. & Aguayo, A. J. Regeneration of long spinal axons in the rat. J Neurocytol 13, 165–82 (1984).

48. Si–Wei You, Kwok–Fai So & Henry K. Yip. Axonal Regeneration of Retinal Ganglion Cells Depending on the Distance of Axotomy in Adult Hamsters | IOVS | ARVO Journals. Investigative Ophthalmology and Visual Sciences 41, 3165–3170 (2000).

49. Jenkins, R., Tetzlaff, W. & Hunt, S. P. Differential Expression of Immediate Early Genes in Rubrospinal Neurons Following Axotomy in Rat. European Journal of Neuroscience 5, 203–209 (1993).

50. Tetzlaff, W., Alexander, S. W., Miller, F. D. & Bisby, M. A. Response of facial and rubrospinal neurons to axotomy: changes in mRNA expression for cytoskeletal proteins and GAP-43. J Neurosci 11, 2528–2544 (1991).

51. Heiman, M., Kulicke, R., Fenster, R. J., Greengard, P. & Heintz, N. Cell type-specific mRNA purification by translating ribosome affinity purification (TRAP). Nat Protoc 9, 1282–1291 (2014).

52. Katz, H. R., Arcese, A. A., Bloom, O. & Morgan, J. R. Activating Transcription Factor 3 (ATF3) is a Highly Conserved Pro-regenerative Transcription Factor in the Vertebrate Nervous System. Front Cell Dev Biol 10, (2022).

53. Shin, J. E., Ha, H., Kim, Y. K., Cho, Y. & DiAntonio, A. DLK regulates a distinctive transcriptional regeneration program after peripheral nerve injury. Neurobiol Dis 127, 178–192 (2019).

54. Asghari Adib, E., Smithson, L. J. & Collins, C. A. An axonal stress response pathway: degenerative and regenerative signaling by DLK. Curr Opin Neurobiol 53, 110–119 (2018).

55. Shin, J. E. et al. Dual Leucine Zipper Kinase Is Required for Retrograde Injury Signaling and Axonal Regeneration. Neuron 74, 1015–1022 (2012).

56. Saikia, J. M. et al. A Critical Role for DLK and LZK in Axonal Repair in the Mammalian Spinal Cord. J Neurosci 42, 3716–3732 (2022).

57. Mahar, M. & Cavalli, V. Intrinsic mechanisms of neuronal axon regeneration. Nature Reviews Neuroscience vol. 19 323–337 Preprint at 10.1038/s41583-018-0001-8 (2018).

58. Sun, F. & He, Z. Neuronal intrinsic barriers for axon regeneration in the adult CNS. Curr Opin Neurobiol 20, 510–8 (2010).

59. Blackmore, M. G. Molecular control of axon growth: insights from comparative gene profiling and high-throughput screening. Int Rev Neurobiol 105, 39–70 (2012).

60. Venkatesh, I. et al. Co-occupancy identifies transcription factor co-operation for axon growth. Nat Commun 12, 2555 (2021).

61. Butler, A., Hoffman, P., Smibert, P., Papalexi, E. & Satija, R. Integrating single-cell transcriptomic data across different conditions, technologies, and species. Nat Biotechnol 36, 411 (2018).

62. Wickham, H. ggplot2. (2016) doi:10.1007/978-3-319-24277-4.

63. Ouyang, J. F., Kamaraj, U. S., Cao, E. Y. & Rackham, O. J. L. ShinyCell: simple and sharable visualization of single-cell gene expression data. Bioinformatics 37, 3374–3376 (2021).

64. Yu, G., Wang, L. G., Han, Y. & He, Q. Y. clusterProfiler: an R Package for Comparing Biological Themes Among Gene Clusters. OMICS 16, 284 (2012).

65. Wu, T. et al. clusterProfiler 4.0: A universal enrichment tool for interpreting omics data. Innovation (Cambridge (Mass.)) 2, (2021).

66. Shannon, P. et al. Cytoscape: A Software Environment for Integrated Models of Biomolecular Interaction Networks. Genome Res 13, 2498 (2003).

67. Wang, Z., Maunze, B., Wang, Y., Tsoulfas, P. & Blackmore, M. G. Global connectivity and function of descending spinal input revealed by 3D microscopy and retrograde transduction. The Journal of Neuroscience 1196–18 (2018) doi:10.1523/JNEUROSCI.1196-18.2018.

68. Soderblom, C. et al. 3D Imaging of Axons in Transparent Spinal Cords from Rodents and Nonhuman Primates. eNeuro 2, (2015).

69. Zheng, G. X. Y. et al. Massively parallel digital transcriptional profiling of single cells. Nature Communications 2017 8:1 8, 1–12 (2017).

