## Supplemental Figures for "Injury distance limits the transcriptional response to spinal injury"

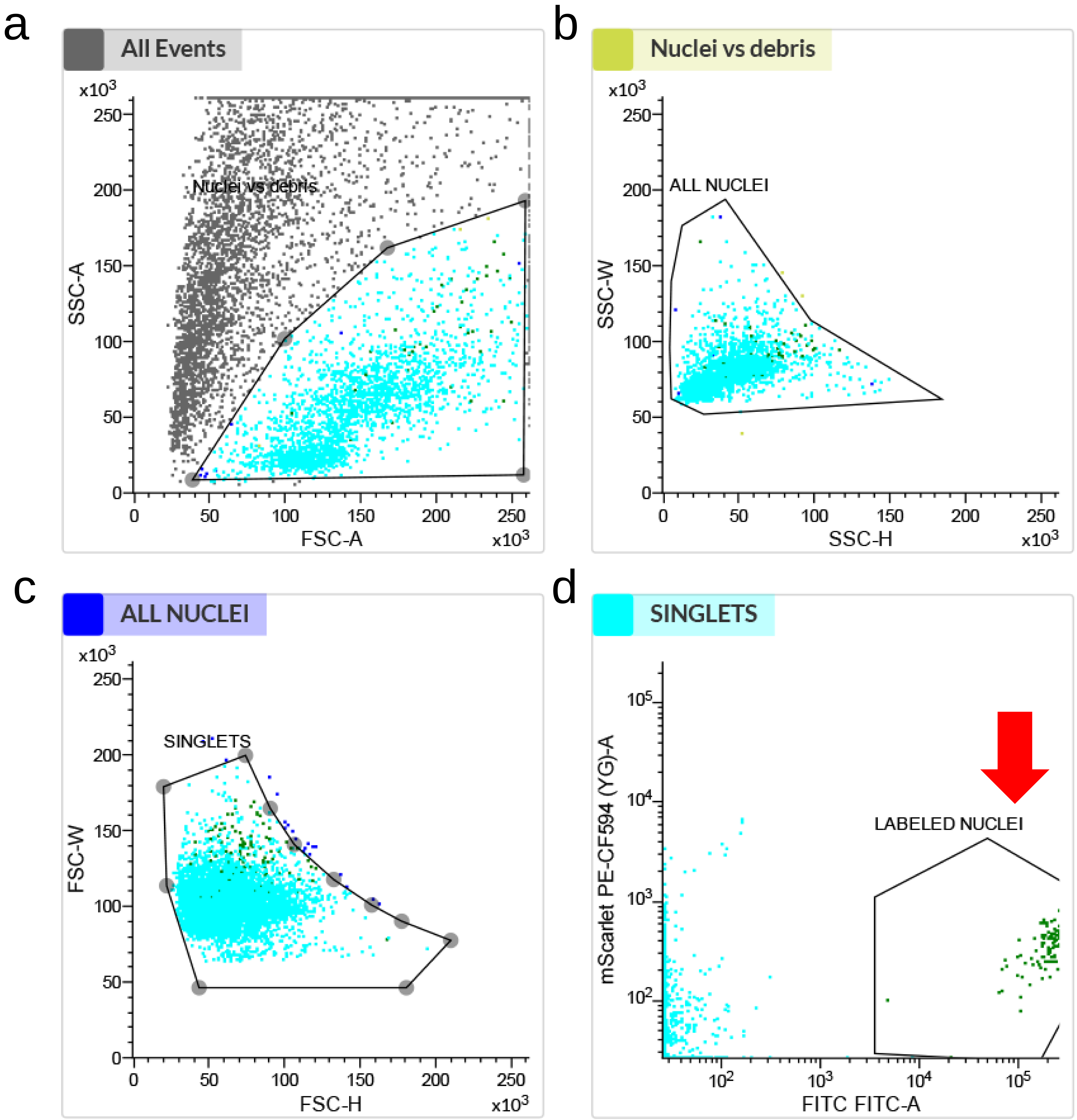


**Supplemental Figure 1: FANS purification of retrogradely labeled supraspinal nuclei.** Nuclei were passed through a series of gates based on forward and side scatter to differentiate nuclei from cellular debris (a, b), minimize doublets (c), and then gather nuclei labeled by mGreenlantern as detected in the FITC channel (d, red arrow).

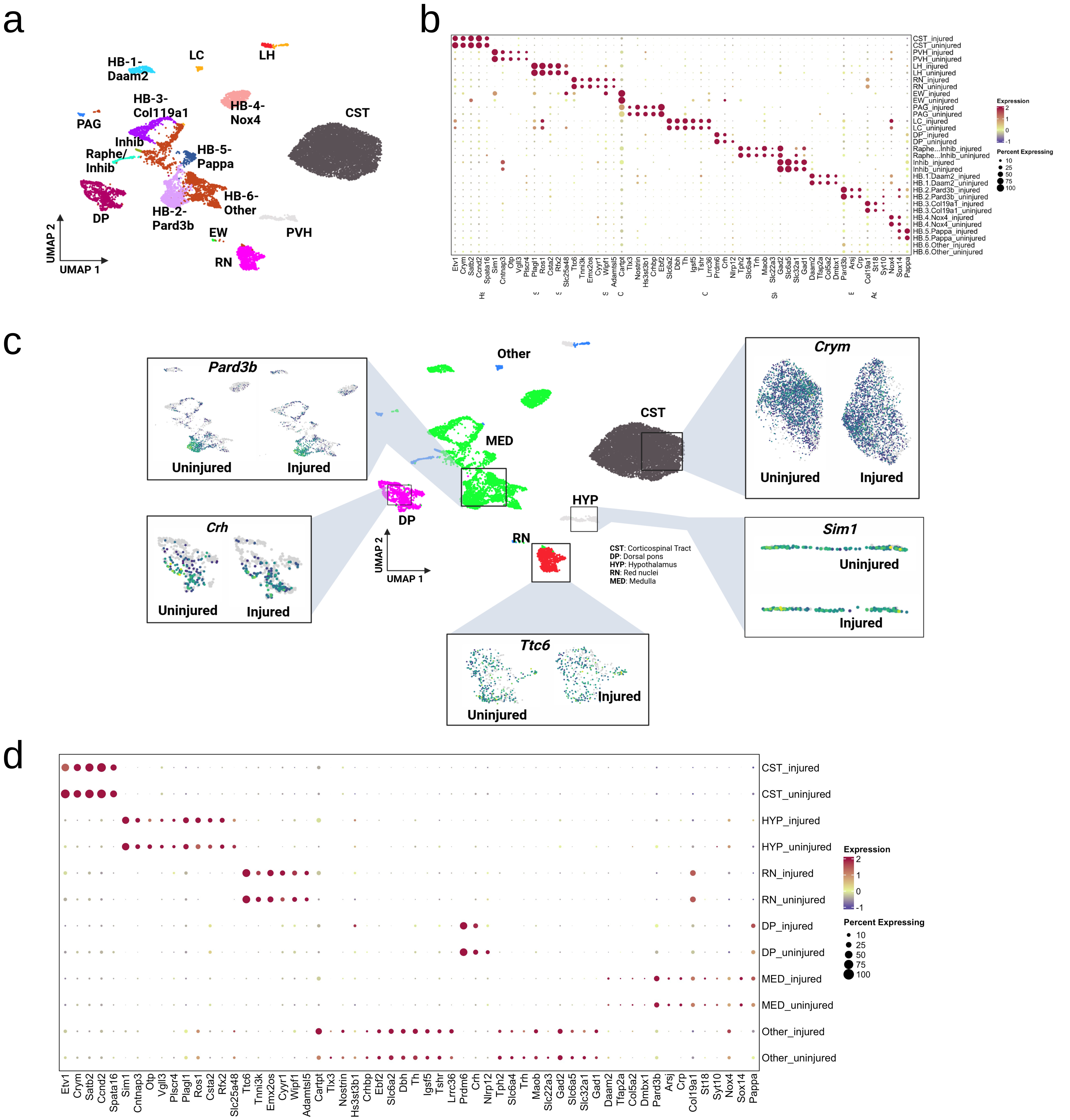


**Supplemental Figure 2: Supraspinal neurons maintain subtype-specific gene expression after thoracic injury.** (a) UMAP clustering of combined injured and uninjured samples shows 16 discrete supra-lumbar cell types . (b) A dot plot shows specific expression of marker genes within the identified clusters and with similar levels of expression in injured and uninjured samples (c) Feature plots illustrate similar levels of marker genes within injured and uninjured samples (d) Regional Expression Dotplot, in which neuronal subtypes are grouped into five main categories: CST (Corticospinal), HYP (Hypothalamic) RN (Red Nucleus), DP (Dorsal Pons), MED (medullary, predominantly reticular), and other. The expression of marker genes is similar in injured an uninjured samples.

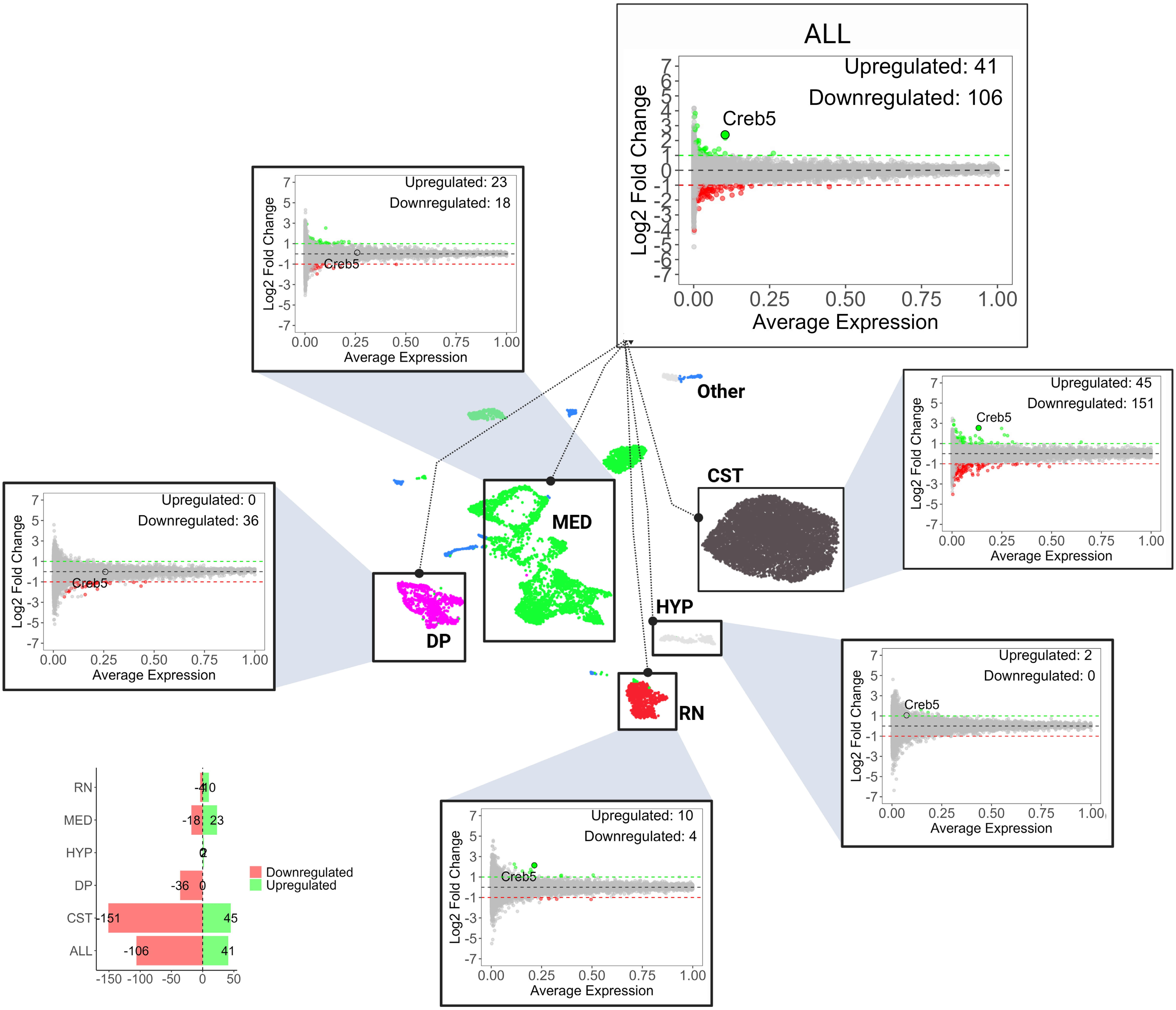


**CST**: Corticospinal Tract **DP**: Dorsal pons **HYP**: Hypothalamus **RN**: Red nuclei **MED**: Medulla

**Supplemental Figure 3. Thoracic injury triggers only modest gene changes in diverse subtypes of supraspinal neurons.** MA plots show changes in gene expression within various brain regions and cell types following thoracic injury (pooled). These regions include the corticospinal tract (CST), dorsal pons (DP), hypothalamus (HYP), red nuclei (RN), and medulla (MED), as well as a composite overview of all regions (ALL). Upregulated genes are marked in green, downregulated genes in red, and the total counts of differentially expressed genes are indicated for each plot, with significance established at a p-value of less than 0.05 via a non-parametric Wilcoxon rank sum test. A bar diagram provides a summary of the counts of differentially expressed genes across the studied regions, with green bars representing the number of upregulated genes and red bars indicating the number of downregulated genes.

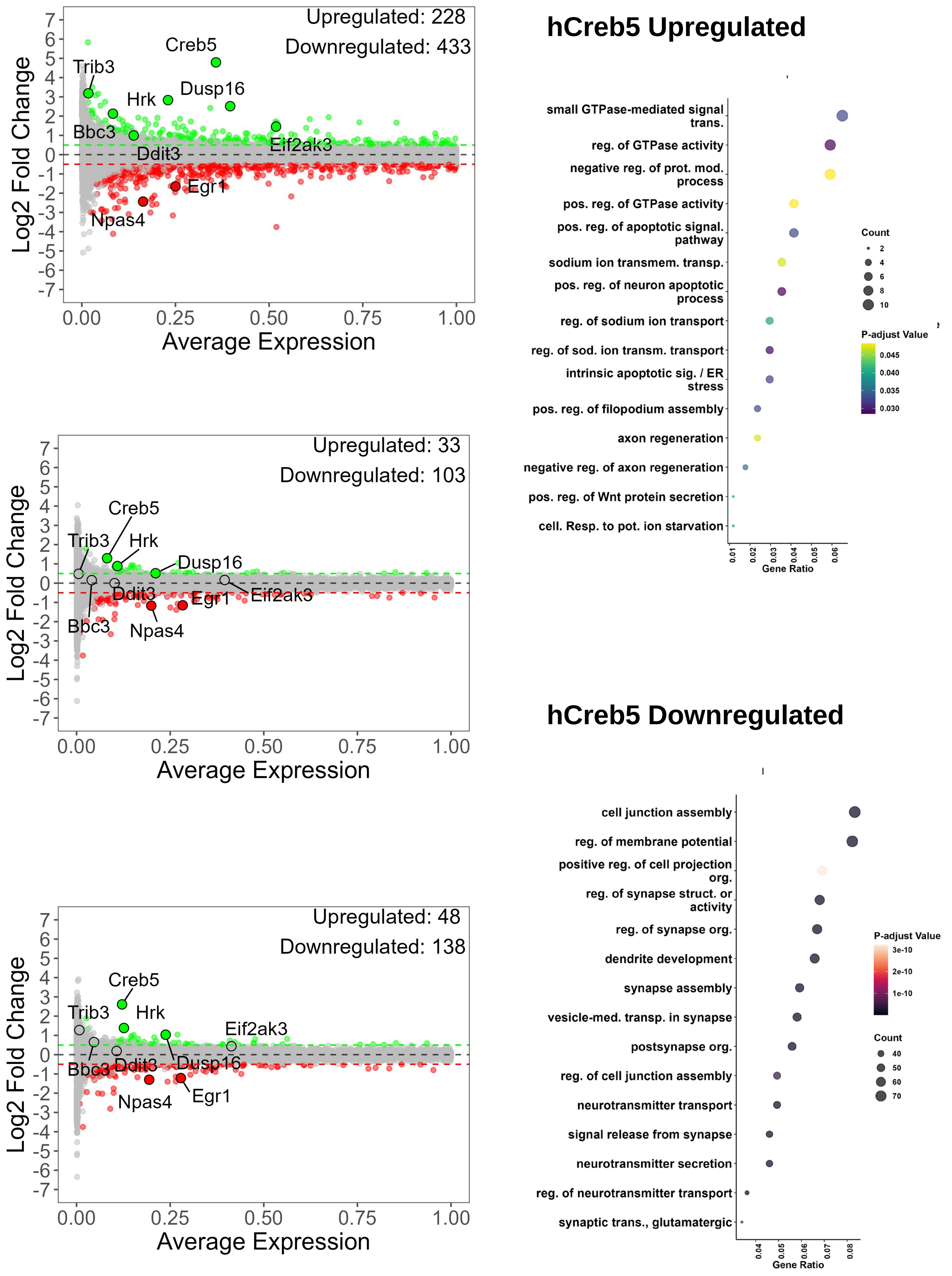


**Supplemental Figure 4: Differential Gene Expression and Pathway Analysis in Corticospinal Tract Neurons after cervical spinal injury.** Injured CST neurons were subdivided according to the level of *Creb5* expression and then compared to uninjured CST neurons. The MA plots alongside reveal differential gene expression, where genes that are upregulated post-injury are marked in green and those that are downregulated in red, with counts for each category provided. Additionally, the log2 fold change in expression after injury relative to uninjured controls is shown (p-value of less than 0.05 via a non-parametric Wilcoxon rank sum test). Gene Ontology (GO) enrichment plots identify enriched GO terms for upregulated and downregulated genes in high-Creb5 CST nuclei. The size of the dots in these plots corresponds to the number of genes associated with each GO term, while the color intensity reflects the significance of enrichment (-log10 adjusted p-value). Within the high-Creb5 subset of CST neurons, upregulated genes are enriched for ER stress, apoptosis, and axon regeneration, while downregulated genes are enriched for synaptic functions.

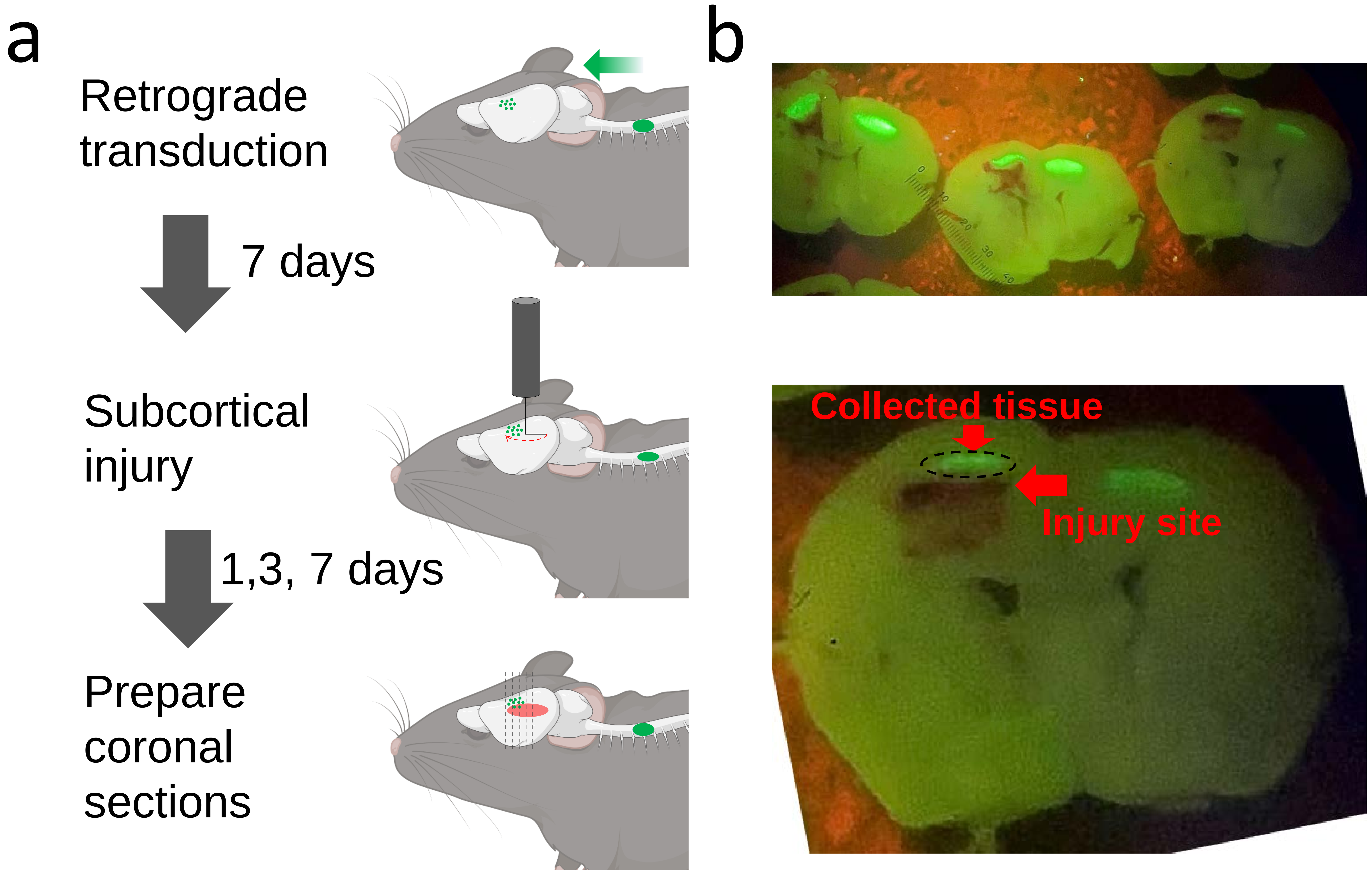


**Supplemental Figure 5: Intracortical axon injury and collection of proximally injured CST neurons.** (a) Experimental approach. CST cell nuclei were retrogradely labeled by cervical injection of AAV2-retro-H2B-mGl. One week later a right-angle blade was lowered into caudal cortex and then rotated 180°, swinging the blade in a path just ventral to CST cell bodies and transecting axons. (b) In coronal sections of brain the CST cell layer is labeled green and the path of the blade through deep cortex is indicated by a line of hemmorage (red arrow). Dotted black line indicates the area of tissue collected for single-nuclei library preparation.

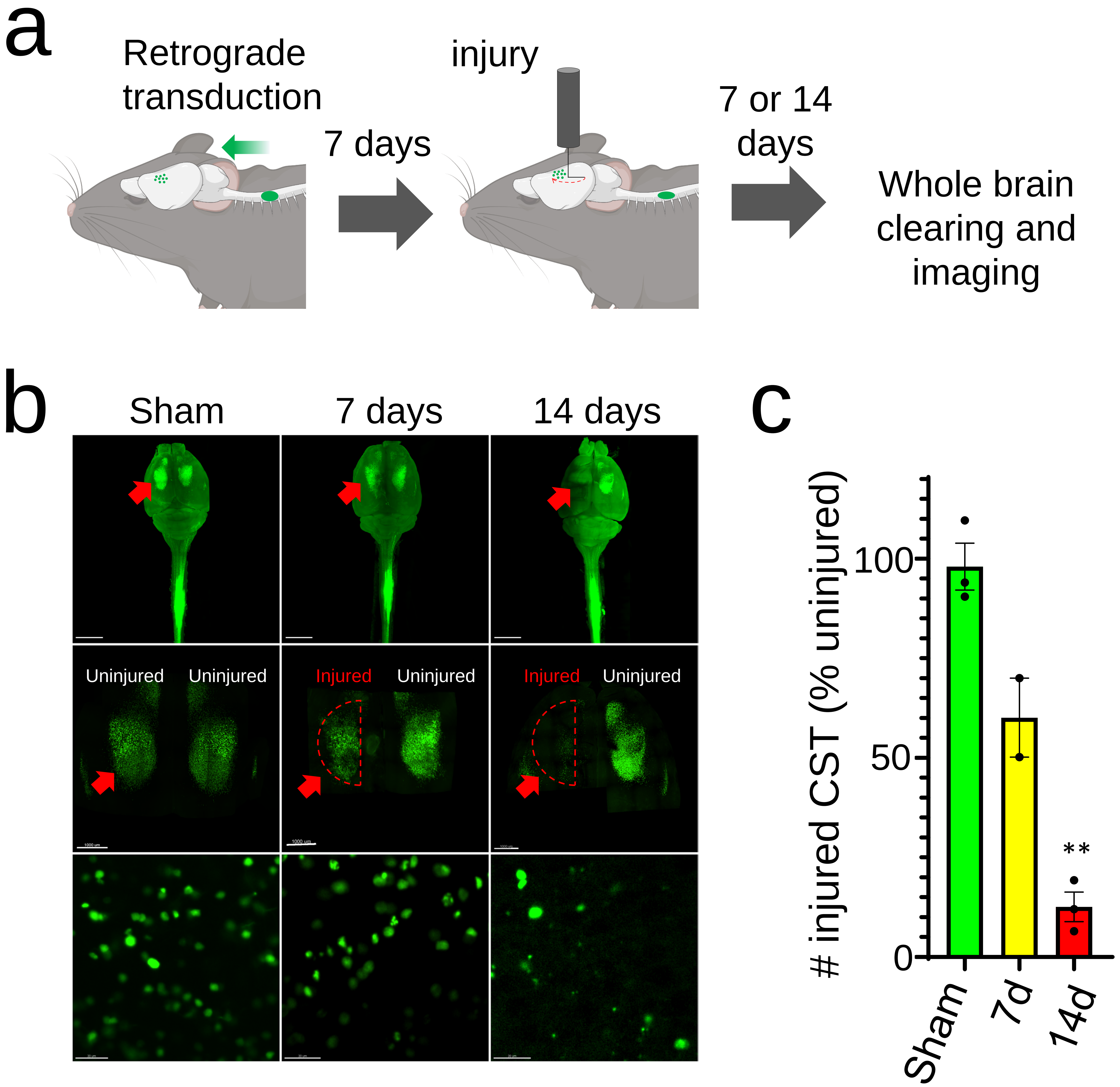


**Supplemental Figure 6: Intracortical axon injury causes loss of CST neurons.**
(a) Experimental design. CST cell nuclei were retrogradely labeled by cervical injection of AAV2-retro-H2B-mGl followed by intracortical injury and whole brain imaging after 7 or 14 days. (b) Dorsal views of transparent cortex with CST cell nuclei labeled in green and approximate injury location shown in red. (c) Quantification of detected CST nuclei overlying the area of cortical injury, normalized to the number detected in the corresponding area in contralateral cortex, reveals a decline of approximately 40% by one week and 85% by two weeks. N=3 animals were time point. **p<.01, ANOVA with post-hoc Sidak's.

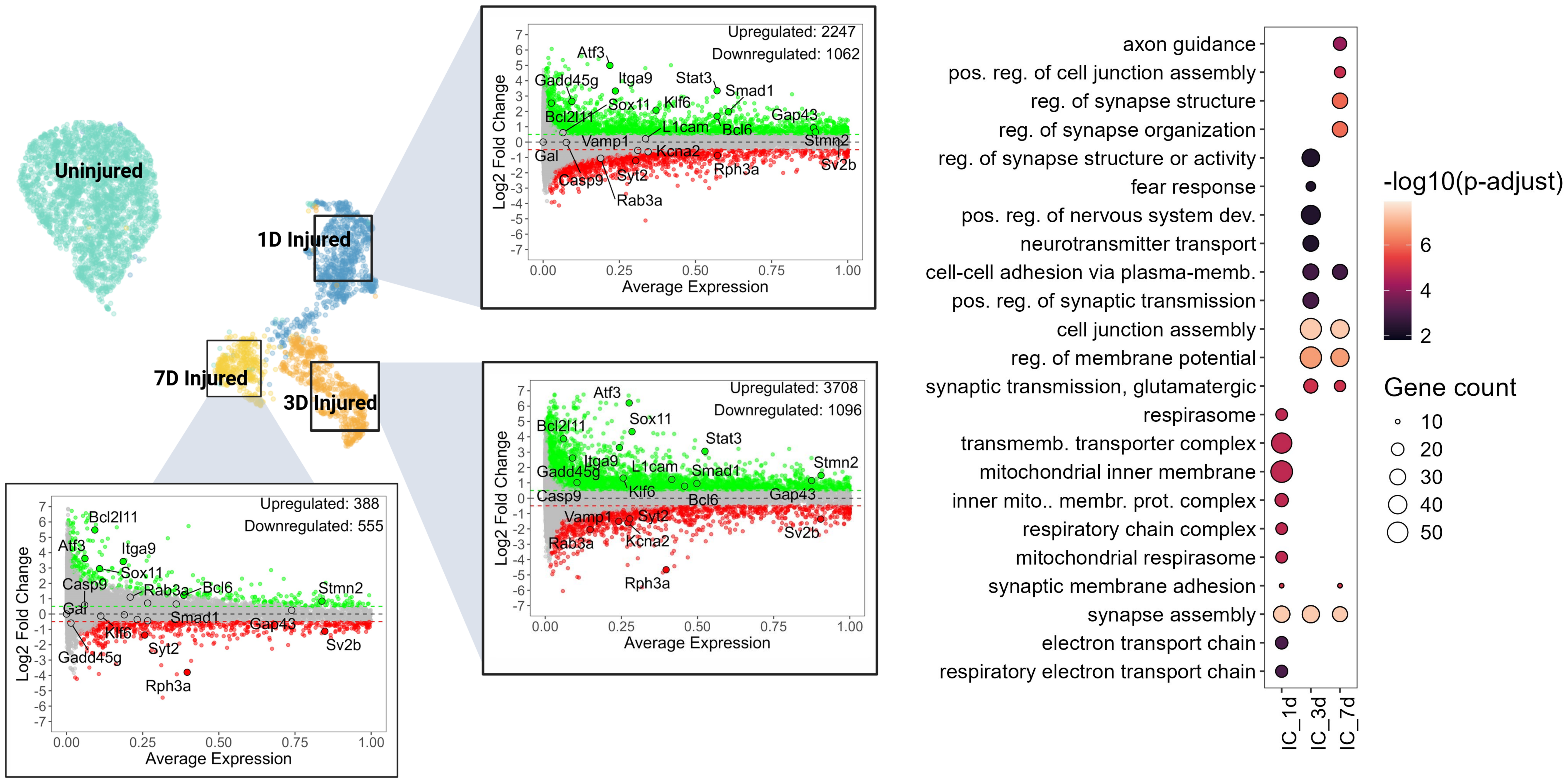


**Supplemental Figure 7: Transcripts downregulated in CST neurons after intracortical injury are enriched for synaptic functions.** UMAP plot categorizes nuclei from both uninjured and injured CST by time point post-injury (teal, 1 day post-injury (DPI) in dark blue, 3 DPI in orange, and 7 DPI in yellow. MA plots display the extent of gene expression downregulation (in red) and upregulation (in green) for each time point (non-parametric Wilcoxon rank sum test, p-value<0.05). A dot plot for Gene Ontology (GO) Enrichment Analysis illustrates the downregulated genes' enriched GO terms over the time points post-injury, spotlighting pathways with fold change values under 0.5. Color intensity indicates the significance level (-log10 of the adjusted p-value), and dot size indicates the number of genes associated with each term.

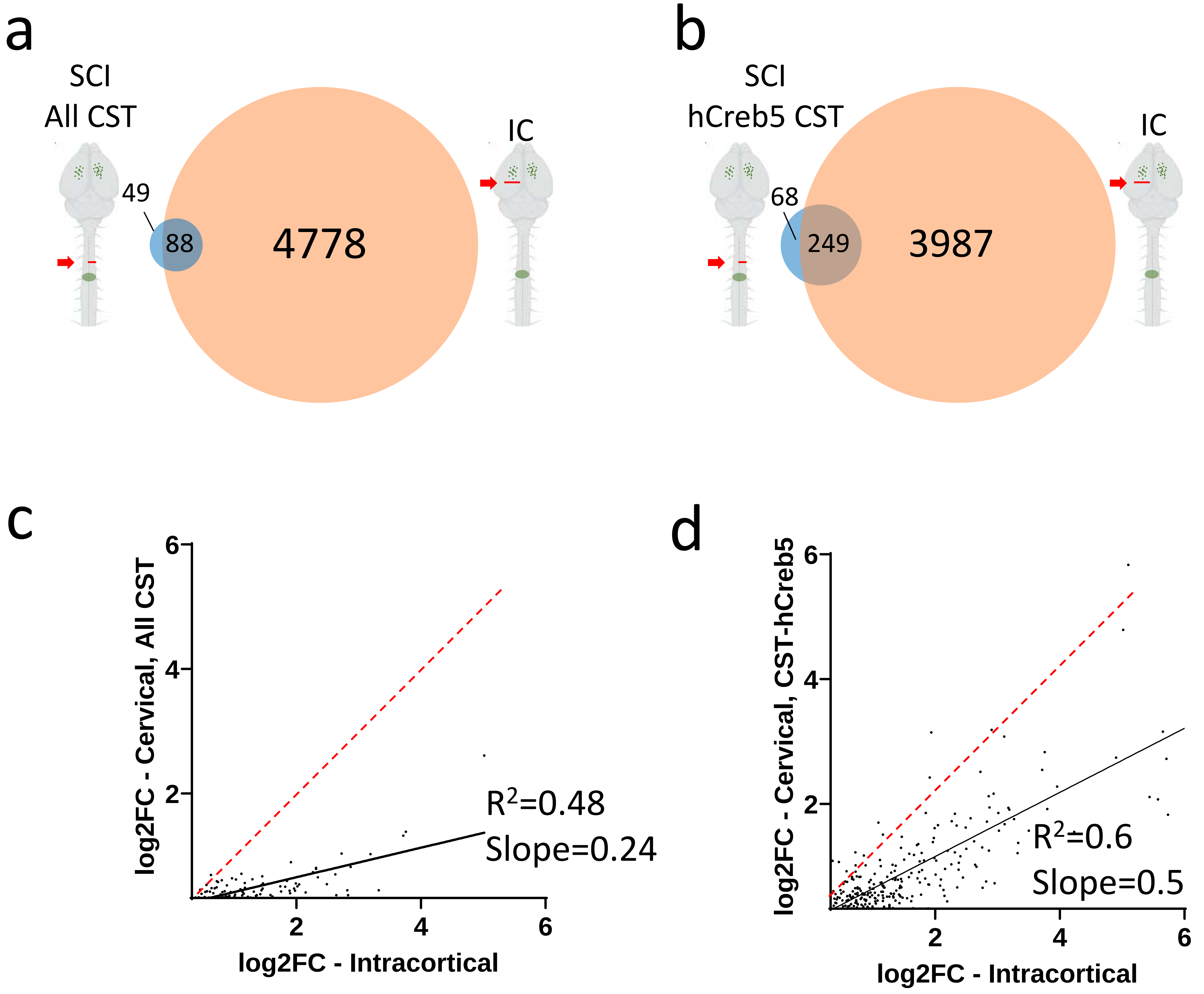


**Supplemental Figure 8: Transcripts upregulated by spinal injury mostly overlap with intracortical-upregulated transcripts but show a lower fold change.** (a) Comparison of transcripts significantly increased more than 25% (log2FC > 0.32) in CST neurons after spinal injury to transcripts increased after intracortical injury shows significant overlap (64%, p<.0001, hypergeometric). (b) A similar comparison using only the stronger-responding subset of cervically injured CST neurons (CST-hCreb5) shows 79% overlap (p<.0001, hypergeometric). (c,d) The fold change values in commonly upregulated transcripts are positively correlated with a slope significantly less than 1 (p<.001, F-test), indicating larger fold changes after intracortical that cervical injury.
